# Evolution of class-structured populations in periodic environments

**DOI:** 10.1101/2021.03.12.435065

**Authors:** Sébastien Lion, Sylvain Gandon

## Abstract

What is the influence of periodic environmental fluctuations on life-history evolution? We present a general theoretical framework to understand and predict the long-term evolution of lifehistory traits under a broad range of ecological scenarios. Specifically, we investigate how periodic fluctuations affect selection when the population is also structured in distinct classes. This analysis yields time-varying selection gradients that clarify the influence of the fluctuations of the environment on the competitive ability of a specific life-history mutation. We use this framework to analyse the evolution of key life-history traits of pathogens. We examine three different epidemiological scenarios and we show how periodic fluctuations of the environment can affect the evolution of virulence and transmission as well as the preference for different hosts. These examples yield new and testable predictions on pathogen evolution, and illustrate how our approach can provide a better understanding of the evolutionary consequences of time-varying environmental fluctuations in a broad range of scenarios.

## 1 Introduction

Many organisms experience periodic fluctuations of their environment. These fluctuations may be driven by abiotic variations of the environment at different time scales (e.g. diurnal and seasonal variability), or by the dynamics of biotic interactions between organisms (e.g. predator-prey or hostparasite limit cycles). In such periodically changing environments, selection on life-history traits is likely to fluctuate over time, but we currently lack a good understanding of the feedback between periodic environmental dynamics and long-term phenotypic evolution (Barraquand et al., 2017).

A good measure of selection in periodic environments should tell us whether, on average over one period of the fluctuation, a mutation increases or decreases in frequency. But how should we compute this average fitness when selection may vary both in time but also among different classes of individuals? Floquet theory provides an answer to this question through the computation of the invasion fitness of a rare mutant in the periodic environment produced by the wild type (Metz et al., 1992; Meszéna et al., 2005; Klausmeier, 2008; Metz, 2008). However, the analysis based on Floquet theory is numerical and yields little biological insight. It only provides a good understanding of evolutionary dynamics when a single class of individuals is needed to describe the mutant dynamics. In this case the invasion fitness is simply the average, over one period, of the per-capita growth rate of a rare mutant (Metz, 2008; Donnelly et al., 2013; Kremer & Klausmeier, 2013; Cornet et al., 2014; Gandon, 2016; Ferris & Best, 2018; Pigeault et al., 2018). In class-structured populations, however, the lack of an analytical expression for the invasion fitness hampers the biological interpretation of the results obtained with Floquet analysis. In this paper, we fill this gap and provide a new method to analyse selection in class-structured populations subject to periodic environmental fluctuations.

In constant environments, it has been shown that the direction of selection should depend on the relative abundance of each class as well as the productivity of the focal organism in each class, so that we need to keep track of both the quantity and the quality of different classes (Taylor, 1990; Taylor & Frank, 1996; Gandon, 2004; Rousset, 2004; Lehmann & Rousset, 2014; Lion, 2018a). Our approach extends this idea to periodic environments and allows us to derive, using only the standard weak-selection assumption, an expression of the selection gradient in terms of the quantity and quality of classes, which are now time-dependent variables (see figure 1 for a graphical summary). With this approach, the selection gradients in periodic and constant environments are directly comparable and conceptually similar.

**Figure 1:**
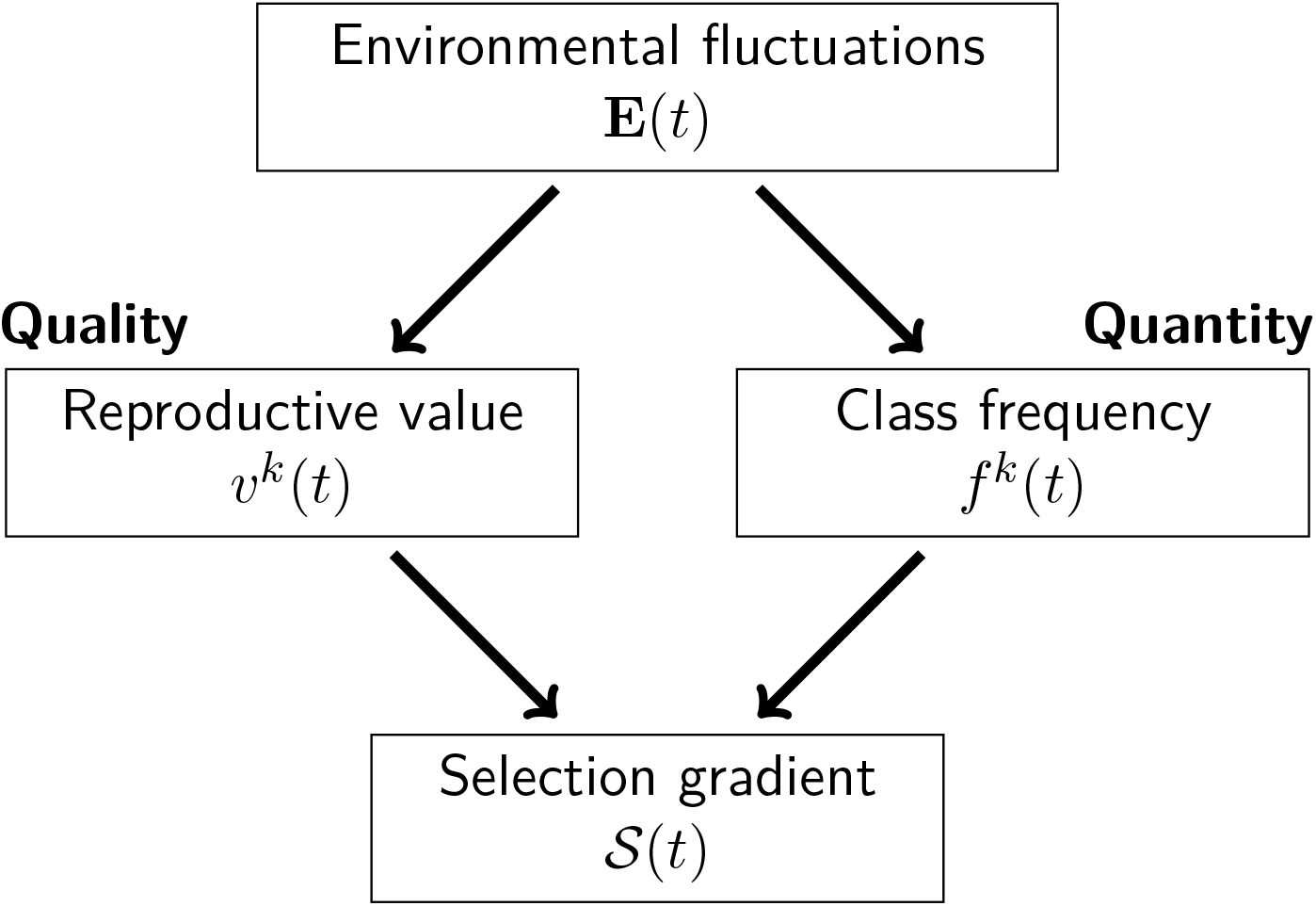
Graphical summary of our approach. Environmental fluctuations are used to evaluate the quality and quantity of individuals in the different classes at a given time. This information can then be used to calculate the selection gradient.

We first provide a general description of eco-evolutionary dynamics in class-structured, polymorphic populations, then turn to the dynamics of a mutant invading a resident population. We show that, under weak selection (that is, for mutations of small phenotypic effects), a separation of time scales argument can be used to derive the selection gradient in periodically varying environments. To illustrate the potential use of this approach, we focus on the evolution of pathogen life-history traits (such as transmission and virulence) in three different epidemiological scenarios when there is periodic variation in the availability of susceptible hosts. The focus on pathogens is not restrictive, and the method can be applied to a variety of life cycles. Evolutionary epidemiology, however, provides a very natural framework in which to think about potentially complex eco-evolutionary feedbacks.

## 2 Eco-evolutionary dynamics

We consider a focal population composed of *K* different *classes* of individuals. For instance, these different classes may correspond to distinct developmental stages of the organism (e.g., young and old, male or female), different immune states, or different locations in a spatially structured environment. Because we are interested in evolution we also assume that the population composed of different *genotypes*.

The life cycle is defined by a matrix of average transition rates, 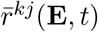, which refer to the net production of class-*k* individuals by class–*j* individuals, averaged over all genotypes (Lion, 2018a,b). These transitions can be due to reproduction, mortality, maturation, or dispersal depending on the biological context. Crucially, these rates can vary with a change in the *environment* which is referred to as **E**(*t*) (Metz et al., 1992; Metz, 2008; Lion, 2018b). These environmental variations may be driven by density-dependent effects caused by changes in population densities ***n***(*t*), by frequencydependent effects caused by changes in the frequencies **p**(*t*) of the different types, but also by changes in extrinsic variables (such as the density of a resource) which we refer to as **e**(*t*), so that **E**(*t*) = (**n**(*t*) **p**(*t*) **e**(*t*))^⊤^. Table 1 gives a summary of the main notations.

**Table 1:** Definition of the main mathematical symbols

| Mathematical symbol | Description |
| --- | --- |
| $n^k(t)$ | Density of individuals in class $k$ ($1 \leq k \leq K$) |
| $n(t) = \sum_k n^k(t)$ | Total density of individuals |
| $f^k(t) = n^k(t)/n(t)$ | Frequency of individuals in class $k$ (with respect to the total population) |
| $v^k(t)$ | Individual reproductive value for class $k$ |
| $\mathbf{n}(t)$ | Vector of class densities $n^k(t)$ |
| $\mathbf{f}(t)$ | Vector of class frequencies $f^k(t)$ |
| $\mathbf{v}(t)$ | Vector of individual reproductive values $v^k(t)$ |
| $\mathbf{e}(t)$ | Vector of external ecological variables |
| $f_i^k(t)$ | Frequency of genotype $i$ within class $k$ ($1 \leq i \leq M$) |
| $\mathbf{p}(t)$ | Vector of genotype frequencies $f_i^k$ (dimension $K \times M$) |
| $f_i(t) = \sum_k f_i^k(t) f^k(t)$ | Global frequency of genotype $i$ |
| $\tilde{f}_i(t) = \sum_k v^k(t) f_i^k(t) f^k(t)$ | Reproductive-value-weighted frequency of genotype $i$ |
| $r_i^{kj}$ | Per-capita transition rate of genotype-$i$ individuals from class $j$ to class $k$ |
| $\bar{r}^{kj} = \sum_i r_i^{kj} f_i^j$ | Average per-capita transition rate at from class $j$ to class $k$ |
| $\bar{\mathbf{R}}$ | Matrix of average per-capita transition rates $\bar{r}^{kj}$. |
| $\mathbf{R}_w$ | Matrix of per-capita transition rates for the resident type $r_w^{kj}$. |

The average transition rates depend on the transition rates of the various genotypes, 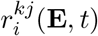, and on the frequencies 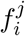 of genotype *i* in class *j*. Thus, we have

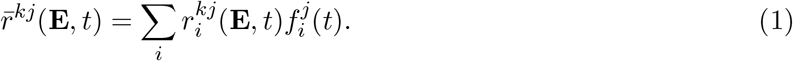

Note that the vital rates may themselves be time-dependent, hence we make time an explicit argument of the transition rates 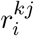. Following Lion (2018a,b), this yields the following eco-evolutionary dynamics:

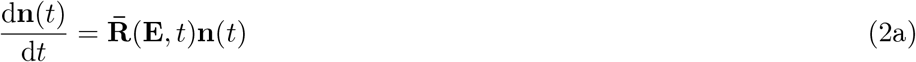

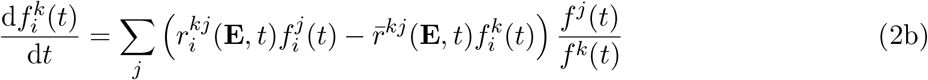

where 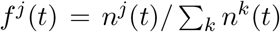 is the fraction of individuals in class *j* at time *t*, and 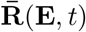 is the matrix of average per-capita transition rates between different classes of individuals. Note that if the environment depends on extrinsic variables (e.g. the density of a resource or a predator), one needs to specify the dynamics of **e**(*t*) to complete the characterisation of the dynamical system (2). Thus, the eco-evolutionary dynamics are described by the *M*(*K* + 1) equations of system (2), plus the equations needed to describe the dynamics of extrinsic variables.

### 2.1 Dynamics of mutant frequencies

Now suppose that, for simplicity, we only have two types in the population: a resident wild type (*w*) and a mutant (*m*). The change in the global frequency of the mutant, 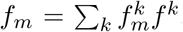, can then be decomposed as follows (Day & Gandon, 2006; Osnas et al., 2015; Lion & Gandon, 2016; Lion, 2018a).

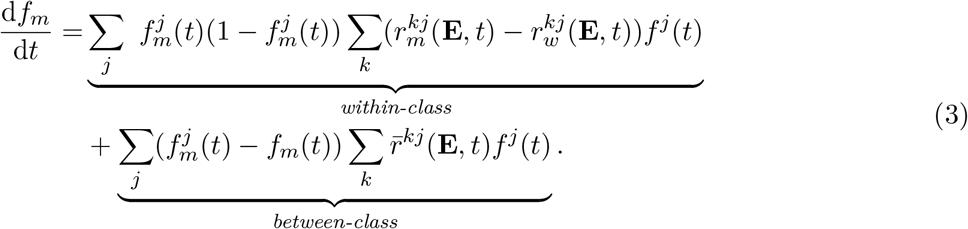

The first line represents the average effect of within-class selection, given by the genetic variance 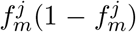 within class *j*, times the difference in transition rates from class *j* to all other classes. The average is taken over the class distribution *f^j^*. In contrast, the second line represents the effect of gene flow between classes, which depends on the relative contribution of the different classes when there is variation in genotype frequencies (differentiation) among classes. Importantly, this second term conflates both the effect of selection (which can shape the differentiation 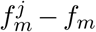 and of purely demographic processes (what Grafen (2015) termed “passive changes” in allele frequencies), so that any estimation of selection based on equation (3) may be biased by the existence of intrinsic differences in qualities between classes (Gardner, 2015; Grafen, 2015; Lion, 2018a).

Thus, although there can be value in explicitly tracking the dynamics of the genetic differentiation between classes (see e.g. Berngruber et al. (2013), Berngruber et al. (2015), and Lion & Gandon (2016)), it is for our purpose more convenient to use an alternative measure of the mutant frequency,

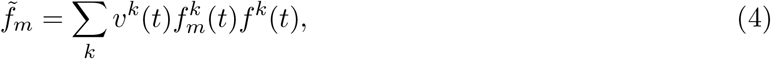

which weights the mutant frequency in class *k* by the *quantity* (*f^k^*(*t*)) and *quality* (*ν^k^*(*t*)) of individuals in class *k* at time *t*. Specifically, we use the reproductive value of an individual in class *k* at time *t* as the measure of quality *ν^k^*(*t*). As previously shown (Lion, 2018a), the dynamics of this weighted average frequency can then be written as

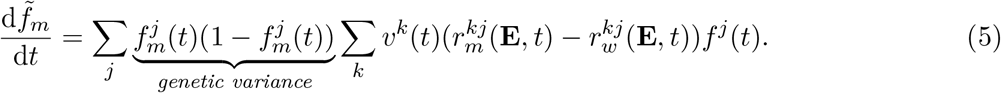

In contrast to equation (3), equation (5) describes the net effect of selection on the change in mutant frequency, without the confounding variations due to “passive changes” (Gardner, 2015; Grafen, 2015; Lion, 2018a). It thus provides a more convenient measure of selection where the overall change in mutant frequency is driven by the sum of the effects of the mutation on the rates *r^kj^* weighted by the frequency of class *j* and the individual reproductive value of class *k*. Crucially, both the quantity and the quality of the different classes can change in a fluctuating environment, so, unlike classical equilibrium theory we need to characterise these fluctuations to understand and predict life-history evolution. In order to characterise these fluctutations, we will use a weak selection assumption to decouple the ecological fluctuations from the evolutionary dynamics.

## 3 Weak selection approximation

In this section, we use a weak-selection approximation of equation (5) to derive an expression of the selection gradient using dynamical reproductive values. Our approach is based on a separation of time scales, which readily occurs when the mutant phenotype, *z_m_*, is close to the resident phenotype *z_w_*, so that *z_m_ = z_w_* + *ε*, where *ε* is small. Under this separation of time scales, the population will settle on a population dynamical attractor, such as a fixed point or a limit cycle, and we can compute the quality and quantity of each class at any given time on the attractor. The fixed point case corresponds to the classical theory developed for constant environments (Taylor, 1990; Rousset, 2004; Otto & Day, 2007), and the limit cycle case corresponds to an extension of this theory to periodic population dynamics, which we now present.

### 3.1 Selection gradient

In appendix A, we show that the class densities **n**(*t*), class frequencies **f**(*t*) and individual reproductive values **v**^⊤^(*t*) are *fast* variables. This means that, under weak selection, the environment **E**(*t*) settles on a periodic attractor which is well approximated by the attractor of a monomorphic resident population, 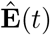. On this monomorphic attractor, we only need the densities 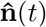 and the external variables 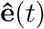 to fully characterise the environment experienced by the mutant type.

In contrast, the weighted mutant frequency 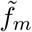 is a *slow* variable. To first-order in *ε*, we can write the dynamics of 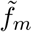 as

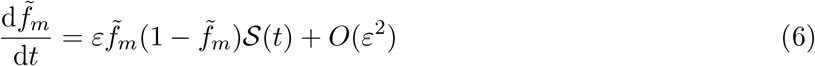

where the instantaneous selection gradient is given by

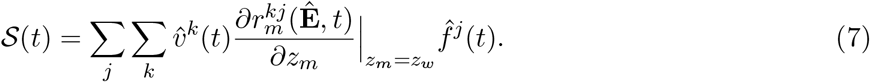

Note that, in equation (6), we have replaced the class variances 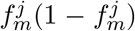 in equation (5) by the population variance 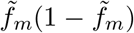. This is because the differences 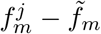 will typically be *O*(*ε*) and therefore this substitution will only contribute an extra *O*(*ε*^2^) term to equation (6).

As a result of periodic fluctuations on the fast time scale, the value and sign of the selection gradient may fluctuate. However, under weak selection, these fast fluctuations can be averaged out. This is known as the averaging principle (see e.g. Cai & Geritz (2020)), which allows us to approximate the dynamics of the mutant frequency on the slow time scale by the solution of the so-called averaged system

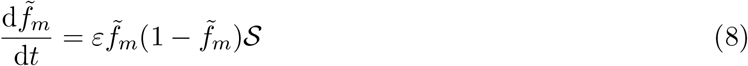

where

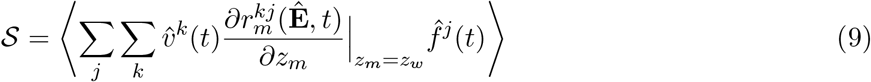

is the selection gradient averaged over one period of the resident attractor (i.e. 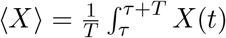 for any given *τ* on the periodic attractor).

The selection gradient (9) takes the form of a sum of marginal selective effects (giving the influence of the evolving trait on between-class transitions) weighted by the time-varying quantity and quality of different classes and is therefore reminiscent of the expressions obtained in equilibrium class-structured populations. In fact, for constant environments, the class frequencies, reproductive values and environment are constant, and therefore equation (9) exactly reduces to the classical expression of the selection gradient for class-structured equilibrium populations (Taylor, 1990; Rousset, 2004; Otto & Day, 2007). For periodic environments, equation (9) provides an analytical approximation, for weak selection, of the invasion fitness of a mutant typically calculated as a Floquet exponent (Appendix B). Note that this first-order approximation gives information on the direction of selection and its potential evolutionary endpoints, but not on the evolutionarily stability of these singularities, for which a numerical computation of the Floquet exponent is still needed (Appendix B).

In equation (9), the class frequencies and reproductive values are calculated on the attractor of the monomorphic resident population. In the next two sections, we show how these quantities can be calculated using dynamical equations of the resident population.

### 3.2 Dynamics of class frequencies

Following Lion (2018a), the class frequencies follow the dynamics:

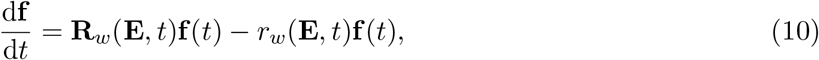

where 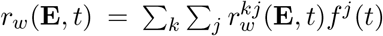 is the per-capita growth rate of the monomorphic resident population and **R**_*w*_ (**E**, *t*) is the matrix of resident transition rates 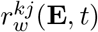. If we focus on the frequency of class *j* we have:

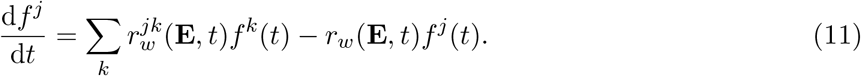

Hence, the change in frequency of class *j* depends on the current class frequencies and on the relative contributions of all the classes to class *j*. Note that we need to account for the overall growth rate *r_w_*(**E**,*t*) of the whole population. This is because we are monitoring the change in class frequencies, not class densities. The sum of class densities can increase (i.e. when *r_w_*(**E**,*t*) > 0) but the sum of class frequencies must remain equal to 1 at all times (i.e. 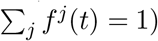).

Mathematically, we can calculate the class frequencies at time *t* by integrating equation (10) forward in time from an appropriate initial condition (i.e. an initial condition that leads to a biologically relevant periodic attractor after some time). For instance, we can use an initial condition where all the *f^j^* are equal to indicate that there is no *a priori* information on the relative quantities of the different classes. In other words, the *quantity* of class *j* at time *t* depends on the *past* trajectory of the population (Figure 2, top panels).

**Figure 2:**
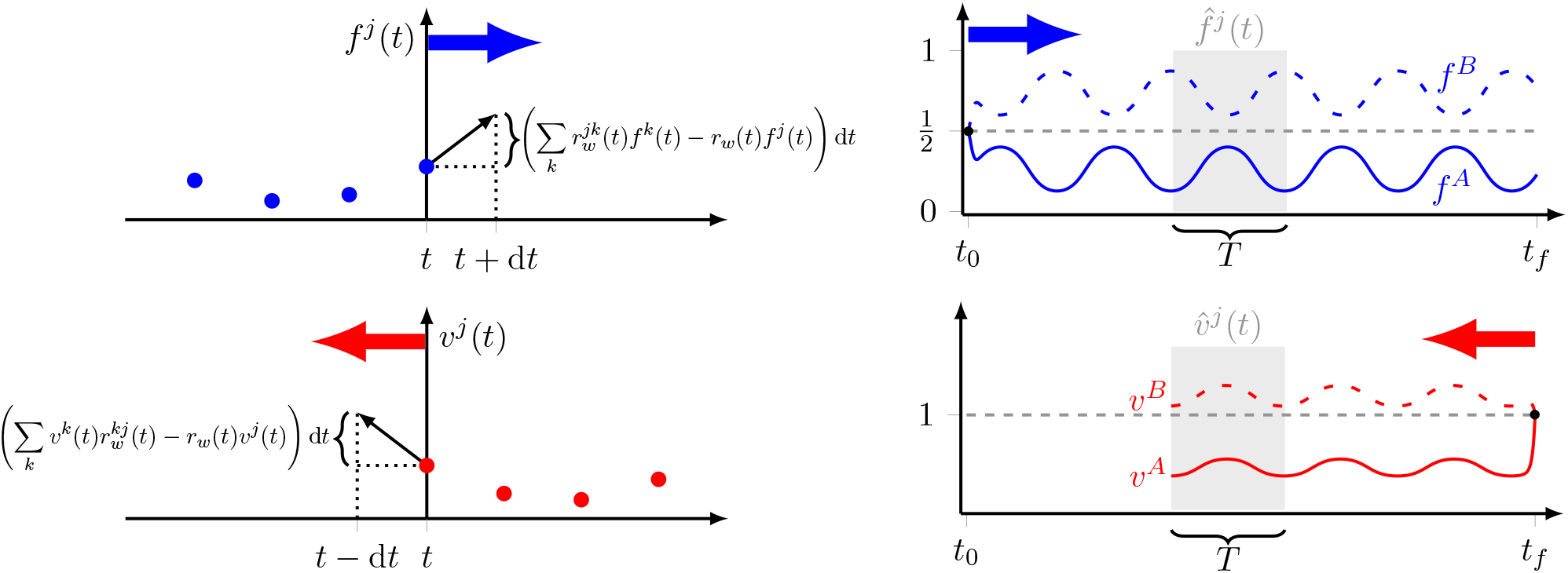
Calculating the class frequencies and reproductive values on the resident periodic attractor. Top panels: The class frequencies are calculated forward in time using equation (11) (left). Integrating from *t* = *t*_0_ to *t_f_*, this gives dynamics that settle on a periodic attractor (right). In this figure, we assume that both classes are initially equifrequent (i.e. *f^A^*(*t*_0_) = *f^B^*(*t*_0_) = 1/2) which amounts to saying that we have no *a priori* information about their relative abundance. – Bottom panels: Next, we insert the values *f^j^* (*t*) into equation (13) to calculate the individual reproductive values. This is a backward process, because we need the values at *t* to compute the values at *t* – d*t* (left). We therefore integrate equation (12) using the final condition *ν^A^*(*t_f_*) = *ν^B^*(*t_f_*) = 1, which means that there is initially (i.e. at time *t_f_*) no *a priori* information on the relative quality of the two classes. We can then choose a period of the attractor (e.g. the gray zone) and calculate the values 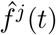 and 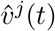 for any *t* in the period.

### 3.3 Dynamics of individual reproductive values

Similarly, the individual reproductive values, collected in the row vector **v**^⊤^(*t*), follow the dynamics (Lion, 2018a):

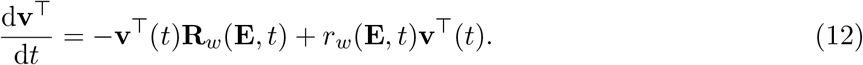

If we focus on the individual reproductive value in class *j*, we have:

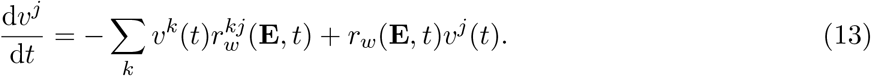

Mathematically, equation (12) is the adjoint of equation (10), which generalises the fact that, in constant environments, class frequencies and individual reproductive values are respectively right and left eigenvectors of the transition matrix of the resident population. Biologically, *ν^k^*(*t*) gives the relative quality of class *k* at time *t*. As in equation (10), we take into account the change in total population size (through *r_w_*(**E**,*t*)) so that **f** and **v**^⊤^ are co-normalised at all times (i.e. **v**^⊤^(*t*)**f**(*t*) = 1). In other words, the reproductive value of an average individual, 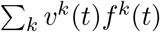 is equal to 1 at all times, and *ν^k^*(*t*) measures the contribution to the future of the population of an individual sampled in class *k* at time *t*, relative to a randomly sampled individual.

Hence, measuring the *quality* of a class at time *t* depends on the descendants and thus on the *future* of the population. In contrast with class frequencies, we compute the individual reproductive values by integrating equation (12) backward in time, from a terminal condition at a time *t_f_* in the distant future (Figure 2, bottom panels). Typically, we set *ν^k^*(*t_f_*) = 1 for all classes at time *t_f_* (Barton & Etheridge, 2011; Lion, 2018a). This terminal condition indicates that there is no *a priori* information on the relative qualities of the different classes of individuals at that point in time and we use between-class transitions on the periodic attractor to acquire information on these relative qualities, by integrating the system (12) backward in time.

### 3.4 A general recipe to study evolution in periodic environments

Equation (9) is the central result of our article and leads to a general recipe to study the evolution of life-history traits in class-structured populations experiencing periodic environmental fluctuations. The main steps of this method are summarised in **Box 1**. We start by computing the dynamics of the class densities in the resident monomorphic population, which allows us to calculate the class frequencies at a given time on the periodic attractor (i.e. in the gray zone in figure 2). We then use this information to calculate the dynamics of the individual reproductive values, which depend on the class frequencies through the computation of the transition rates 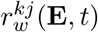. Although in general these solutions can only be computed numerically, we show in the next section that algebraic manipulations of equations (9), (11) and (13) can be used to shed light on the interplay between fluctuating environmental dynamics and selection. We do so by applying our general method to the evolution of pathogen life-history traits when the density of hosts fluctuates.

#### Box 1: How to study life-history evolution in periodically fluctuating environments?

We detail below the different steps allowing us to derive the selection gradient driving life-history evolution in a class-structured population in a periodic environment.

**Step 1 -** Formalise the description of the life cycle of the focal organism in a monomorphic resident population. This is captured in the dynamical system:

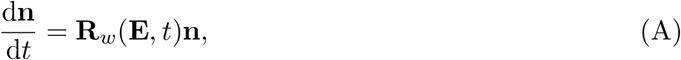

where **R**_*w*_(**E**,*t*) is the matrix of between-class transition rates in the resident population, together with the dynamical equations for the extrinsic variables in **e**(*t*).

**Step 2 -** Determine which life-history traits are under selection (and the trade-off between these traits). Indeed, multiple life-history traits may be involved in the life-cycle and it is crucial to be very explicit about the constraints acting on the evolving traits.

**Step 3 -** Derive the selection gradient using equation (9). This step may often yield insight through the decomposition of selection into biologically meaningful quantities (see section 4 for examples). But in general, moving to steps 4 and 5 is needed to predict the influence of periodic fluctuations on life-history evolution.

**Step 4 -** Use equation (A) and (11) to determine the periodic attractor of the resident population. This will yield the forward dynamics of class frequencies.

**Step 5 -** Use equation (12) and the numerical solution derived in step 4 to solve the backward dynamics of individual reproductive values (using a final condition where individual reproductive values are all equal to 1).

**Step 6 -** The results of steps 4 and 5 can then be plugged into the selection gradient (step 3) to obtain a mathematical expression (or a numerical computation) of the selection gradient on the trait. This can be used to identify evolutionary singularities, the evolutionarily stability of which can be checked using Floquet analysis (Appendix B).

## 4 Pathogen evolution in periodic environments

Many pathogens have to cope with environmental fluctuations (Altizer et al., 2006; Martinez, 2018). In this section, we use the approach in **Box 1** to study pathogen evolution under three distinct epidemiological scenarios corresponding to three different pathogen life cycles. In all scenarios the pathogen can be present in two distinct classes of hosts (*A* and *B*) and the environmental fluctuations are captured by a periodic function *ν*(*t*) that gives the probability of production of susceptible hosts at time *t*. Hence, using our general terminology, the forcing function *ν*(*t*) causes periodic fluctuations in the vector **e**(*t*) (collecting the densities of susceptible hosts), which in turn drives fluctuations in the vector **n**(*t*) (collecting the densities of infected hosts in classes *A* and *B*). The general approach outlined in **Box 1** can be applied to any periodic function, but for simplicity we consider a smooth version of a step function with minimum 0, maximum *ν_max_*, and mean *ν*_*max*/2_ (figure 3).

**Figure 3:**
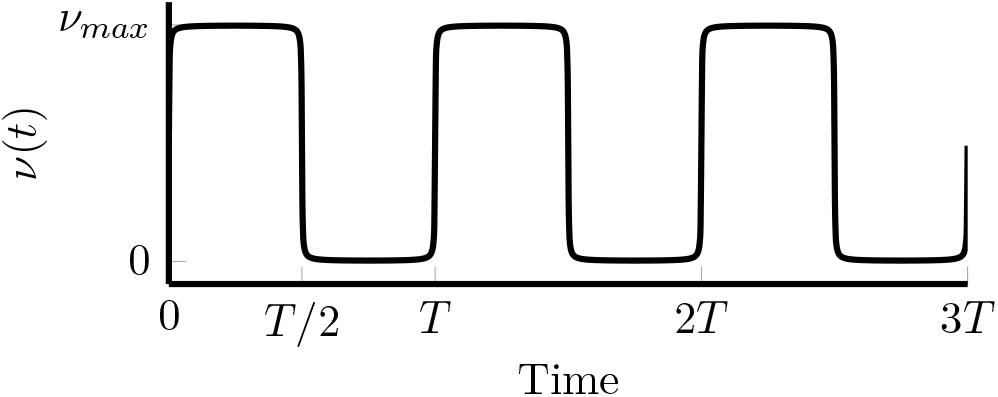
Environmental fluctuation. We model fluctuations in the environment with a smoothed step function with minimum 0, maximum *ν_max_* and mean *ν_max_*/2. This is captured by the following function: *ν*(*t*) = (*ν_max_*/2)(1 + (2*ζ*/*π*) arctan (sin (2*πt*/*T*)/*δ*). We typically use *ζ* = 1 and *δ* = 0.01. Note that *ζ* = 0 corresponds to a constant environment with value *ν* = *ν_max_*/2.

### 4.1 Scenario 1: the Curse of the Pharaoh hypothesis

The claim that long-lived pathogen propagules could select for higher pathogen virulence has often been presented as the “Curse of the Pharaoh hypothesis” (Bonhoeffer et al., 1996; Gandon, 1998). Previous theoretical analyses focused mainly on temporally constant environments. Here we want to analyse the influence of fluctuations on the availability of susceptible hosts on the evolution of virulence for pathogens with free-living stages.

Let us assume that there is a single class of susceptible hosts *S* but two different classes of the pathogens, which are (i) the infected host (*I^A^*) and (ii) the propagule stage (which lives outside the infected host but that we still denote *I^B^* for consistency with the general framework). Thus, the vector of population densities is **n**(*t*) = (*I^A^*(*t*) *I^B^*(*t*) and *S*(*t*) is the extrinsic environmental variable **e**(*t*). The epidemiological dynamics are given by

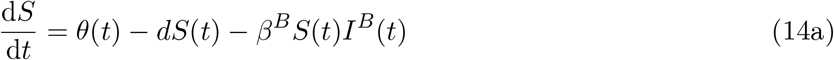

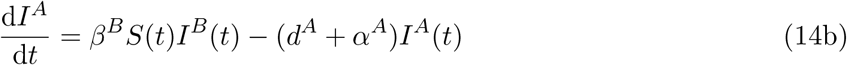

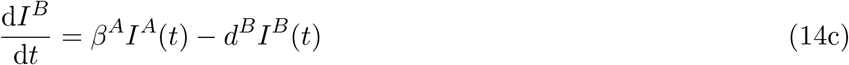

where *θ*(*t*) = *bν*(*t*) refers to the periodic influx of susceptible hosts in the population. A susceptible host becomes infected upon contact with the propagules, which occurs at rate *β^B^I^B^*(*t*). Infected hosts die at rate *d^A^* + *α^A^*, where *d^A^* is the background mortality rate and *α^A^* represents virulence, and produce propagules at rate *β^A^*. These propagules die at a rate *d^B^*. Next, we follow the steps presented in **Box 1** to explore how higher propagule survival (e.g. lower values of *d^B^*) can affect the evolution of pathogen virulence *α^A^* in the infected host.

**Step 1:** We can use equations (14) to derive the matrix **R**_*w*_(**E**, *t*) which captures the transition rates between classes *A* and *B* in a monomorphic resident population:

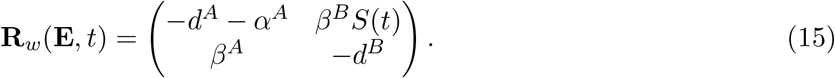

**Step 2:** At this stage it is important to specify the constraints acting on the underlying trait *z* that controls the evolution of pathogen virulence *α^A^*. As in previous studies, we assume that an increase in virulence is associated with an increase in pathogen transmission rate and in particular on the production of propagules *β^A^* (so that the derivatives of *β^A^* and *α^A^* with respect to *z* are both positive; Bonhoeffer et al. (1996) and Day & Gandon (2006)).

**Step 3:** We use equation (7) to obtain the instantaneous selection gradient on the trait *z*:

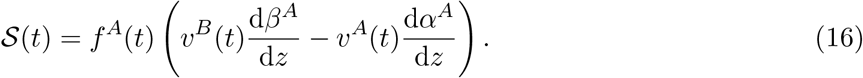

The first term between brackets represents the gain in fitness if a pathogen in class *A* invests in the production of propagules (weighted by the individual reproductive value *ν^B^*(*t*) of propagules). The second term between brackets accounts for the loss in fitness if a pathogen in class *A* dies (weighted by the individual reproductive value *ν^A^*(*t*)). The selection gradient on *z* is obtained by integrating equation (16) over one period of the fluctuation which yields:

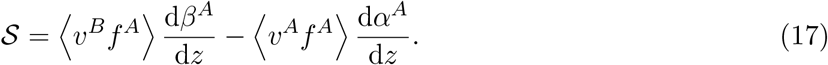

An analysis of the dynamics of class frequencies **f** (*t*) and individual reproductive values **v**(*t*) is required to better understand selection on virulence and transmission.

**Steps 4 and 5:** For our life cycle, equation (12), which gives the dynamics of reproductive values, can be written as:

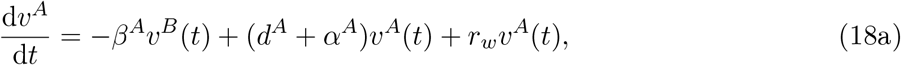

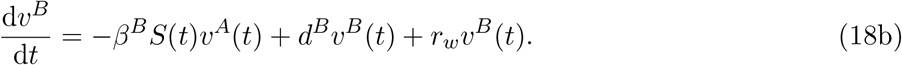

Fluctuations in the availability of susceptible hosts cause fluctuations of the reproductive values which can be obtained numerically from (18), using the solution of system (14) to evaluate *S*(*t*) and 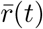. But equation (18) also yields a useful expression for the average of the ratio of individual reproductive values (Appendix S.1):

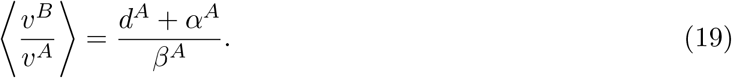

The right-hand side of equation (19) is the ratio of individual reproductive values in the absence of fluctuations, so that we see that the ratio *ν^B^*(*t*)/*ν^A^*(*t*) fluctuates around its equilibrium value in a constant environment.

**Step 6:** Equation (19) can be used to rewrite the selection gradient (17) which yields:

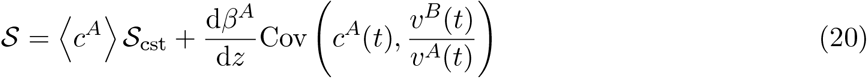

where *c^A^*(*t*) = *ν^A^*(*t*)*f^A^*(*t*) is the class reproductive value at time *t* and

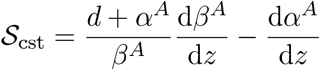

refers to the gradient of selection in the absence of fluctuations. This expression shows that, as pointed out by earlier studies (Bonhoeffer et al., 1996; Day & Gandon, 2006), the mortality rate of the propagule has no effect on the evolutionary stable virulence in a constant environment. However, with periodic fluctuations, we will see that the mortality rate of the propagule does affect the evolutionary outcome. Equation (20) is particularly insightful because it shows how periodic fluctuations of the environment can affect the evolution of the pathogen: because 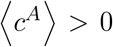, periodic fluctuations will affect the direction of selection only if the temporal covariance between *c^A^*(*t*) and 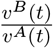 is non-zero. In words, this means that, if infected hosts tend to be more abundant at times when propagules are relatively more valuable (e.g. the covariance is positive), the ES virulence will be higher than in a constant environment because the pathogen then reaps fitness benefits from increased propagule production.

Numerical simulations show that this covariance is expected to be positive when *d^B^* is low and negative when *d^B^* is large (figures 4a, 4b, 4d; see the SOM for an attempt to understand the sign of this covariance). In contrast with the analysis of Bonhoeffer et al. (1996) we thus expect fluctuations to alter the predictions regarding the influence of *d^B^* on the evolution of virulence and transmission rates. As shown in figure 4c, higher rates of survival (i.e. lower values of *d^B^*) tend to select for higher virulence and transmission (as in the Curse of the Pharaoh hypothesis). However, this effect is non-monotonic: when *d^B^* gets very low (i.e. when propagule live very long), the covariance vanishes because *f^A^* → 0 and thus *c^A^* → 0, and therefore the ES virulence is the same as predicted in a constant environment. Note that Figure 4c also shows that, as expected, the ESS predicted from the selection gradient using time-dependent reproductive values is consistent with the value predicted from a more standard Floquet analysis (dashed line).

**Figure 4:**
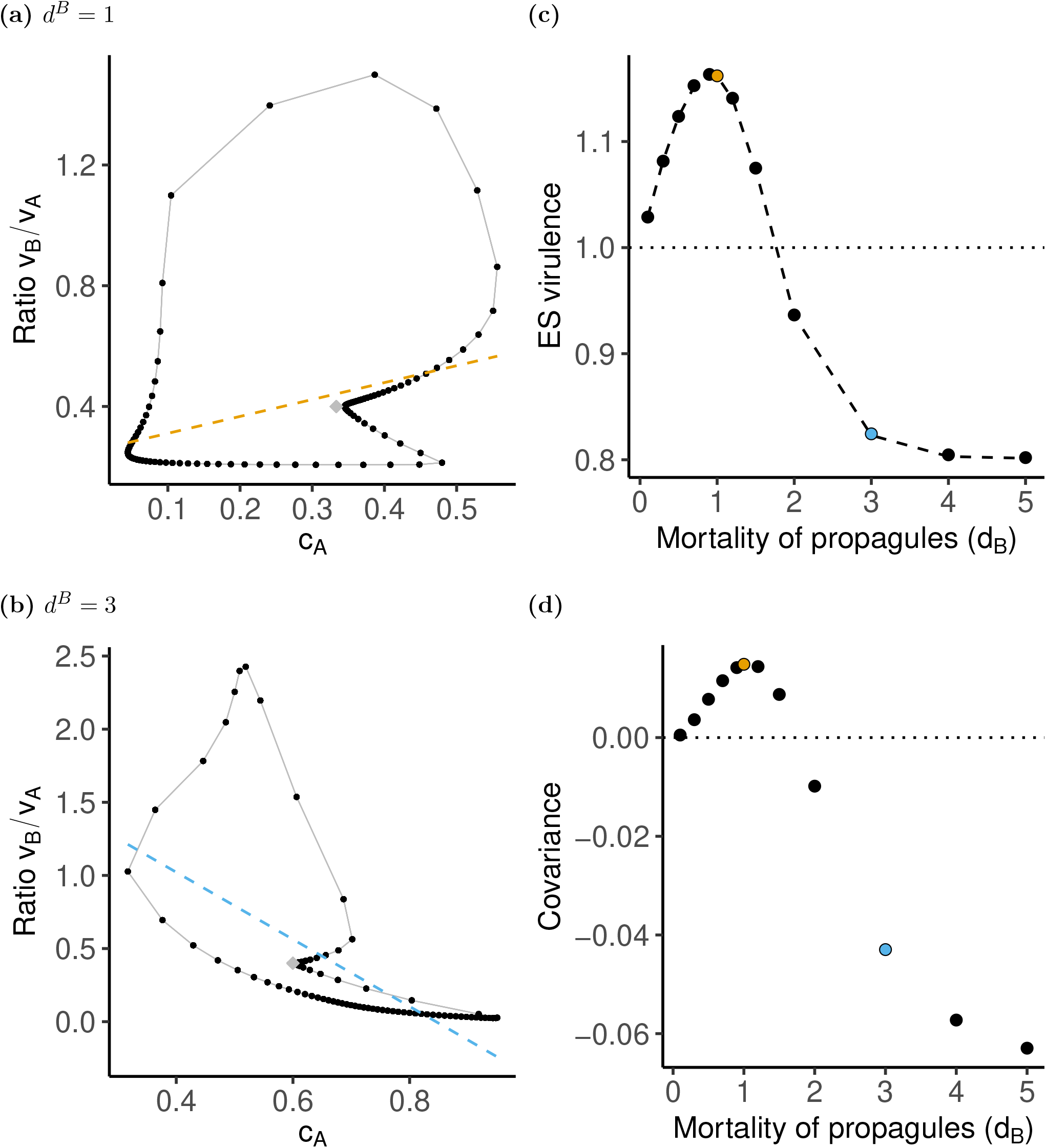
Scenario 1: The Curse of Pharaoh. (a) Parametric plot of *ν^B^*(*t*)/*ν^A^*(*t*) versus *c^A^*(*t*) for *d^B^* = 1 and *z_w_* = *z_eq_*. The grey diamond gives the equilibrium for a constant environment. The slope of the regression line (dashed) is proportional to the temporal covariance between *c^A^*(*t*) and *ν^B^*(*t*)/*ν^A^*(*t*). (b) Same as (a) but with *d^B^* = 3. (c) Predicted ESS as a function of the mortality rate of propagules, *d^B^* using the reproductive-value-based approach (dots) and the Floquet exponent (dashed line). The dotted line gives the prediction of the corresponding equilibrium model, *z_eq_* = 1. (d) Temporal covariance between the class reproductive value *c^A^*(*t*) and the ratio of individual reproductive values *ν^B^*(*t*)/*ν^A^*(*t*) as a function of *d^B^* for *z_w_* = *z_eq_*, where *z_eq_* is the ESS in the absence of fluctuations. Parameters: *ν*(*t*) = 0.5(1 + (2/*π*) arctan (sin (2*πt*/*T*)/0.01)), *b* = 8, *d* = *d^A^* = 1, *β^A^*(*z*) = *β*_0_*z*/(1 + *z*), *β^B^* = *β*_0_ = 10, *α^A^*(*z*) = *z*, *T* = 10.

**Conclusion of Scenario 1:** In a constant environment the longevity of pathogen propagules does not affect the long-term evolution of pathogen virulence. In contrast, we show that periodic fluctuations of the environment (i.e., fluctuations in the availability of susceptible hosts) can select for higher (when the fluctuations are fast) or lower (when the fluctuations are slow) pathogen virulence. The selection gradient given in equation (20) allows us to capture the effect of fluctuations through a single temporal covariance which measures the deviation from the selection gradient in a constant environment.

### 4.2 Scenario 2: host preference

Many pathogens can infect several host species. Intuitively, which host the pathogen should prefer will depend on the relative qualities of the hosts. But what would be the best strategy when the qualities or abundances of the different host species fluctuate? In our second scenario, we investigate the effect of periodic fluctuations in the abundance of different host species on the evolution of the pathogen’s preference strategy.

We consider an epidemiological model with contact transmission (no free-living propagules) and assume that the pathogen can exploit two different hosts (*A* or *B*). When a pathogen enters a susceptible host *S^A^* (resp. *S^B^*), we assume that the infection is successful with probability *p^A^* (resp. *p^B^*). With these assumptions, the dynamical system becomes:

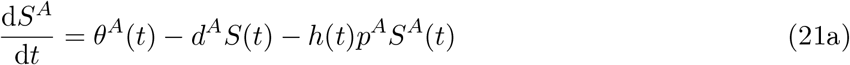

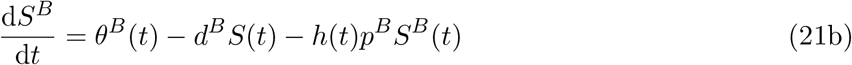

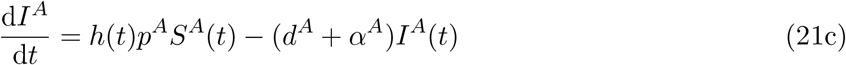

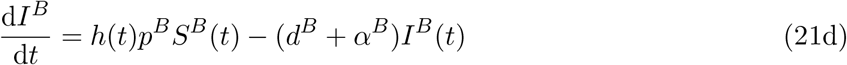

where *h*(*t*) = *β^A^I^A^*(*t*) + *β^B^I^B^*(*t*) is the force of infection. The two infection routes differ by their epidemiological parameters, so that one infection route may be more contagious or virulent than the other. Furthermore, we assume that the production of susceptible hosts is periodic, and such that *θ^A^*(*t*) = *b*(1 – *ν*(*t*)) and *θ^B^* = *bν*(*t*), where *ν*(*t*) is the probability of production of *B* hosts (figure 3).

**Step 1:** From equations (21), we derive the transition matrix

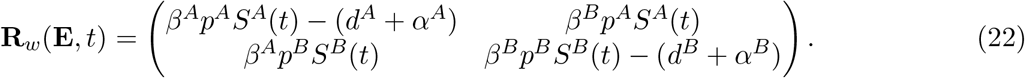

**Step 2:** We assume that there is a trade-off between *p^A^* and *p^B^* such that *p^A^* = *z* = 1 – *p^B^*. Hence, if *z* = 1, infection is only possible on *A* hosts, while if *z* = 1/2, both host classes are equally susceptible to infection. The trait *z* can thus be interpreted as measuring preference towards *A* hosts. For simplicity, we assume that the pathogen’s virulence is lower in host *A*, but its transmissibility is independent of the host (i.e. *α^A^* > *α^B^*, but *β^A^* = *β^B^* = *β*).

**Step 3:** Based on these assumptions on the life cycle, a naive prediction could be that the pathogen should always prefer the “good” host *B*, in which it enjoys a longer lifespan because the pathogen is less virulent on this host. However, the optimal strategy depends on the relative availability of the two classes of hosts, which can fluctuate over time. To better understand the selective pressures on the preference trait, we use our approach to derive the selection gradient at time *t* and obtain:

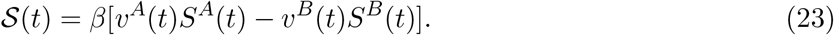

The terms *ν^k^*(*t*)*S^k^*(*t*) have a simple interpretation as the expected reproductive output of a pathogen propagule at time *t* through class *k*. Thus, the direction of selection is determined by whether this reproductive output is larger through class *A* than through class *B*. Furthermore, potential evolutionary endpoints satisfy the balance condition

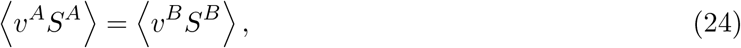

which simply states that selection halts when average reproductive outputs are the same in both classes.

**Steps 4 and 5:** To better understand the impact of periodic fluctuations in host availability on the evolution of the preference trait, we now need to numerically calculate the dynamics of the reproductive values and densities of susceptible hosts. A detailed discussion of these dynamics is given in the SOM, but we show in figures 5a-5b that the period of *ν*(*t*) has a strong impact on the dynamics of the difference in reproductive output *D*(*t*) = *ν^A^*(*t*)*S^A^*(*t*) – *ν^B^*(*t*)*S^B^*(*t*). When the period is small (figure 5a), *D* fluctuates rapidly around a mean that is close to its value in a constant environment (which is zero). In contrast, for large periods (figure 5b), *D*(*t*) better tracks the environmental fluctuations *ν*(*t*) (it is minimal when there are only *B* hosts and maximal when there are only *A* hosts) and its mean 〈*D*〉 is greater than zero.

**Figure 5:**
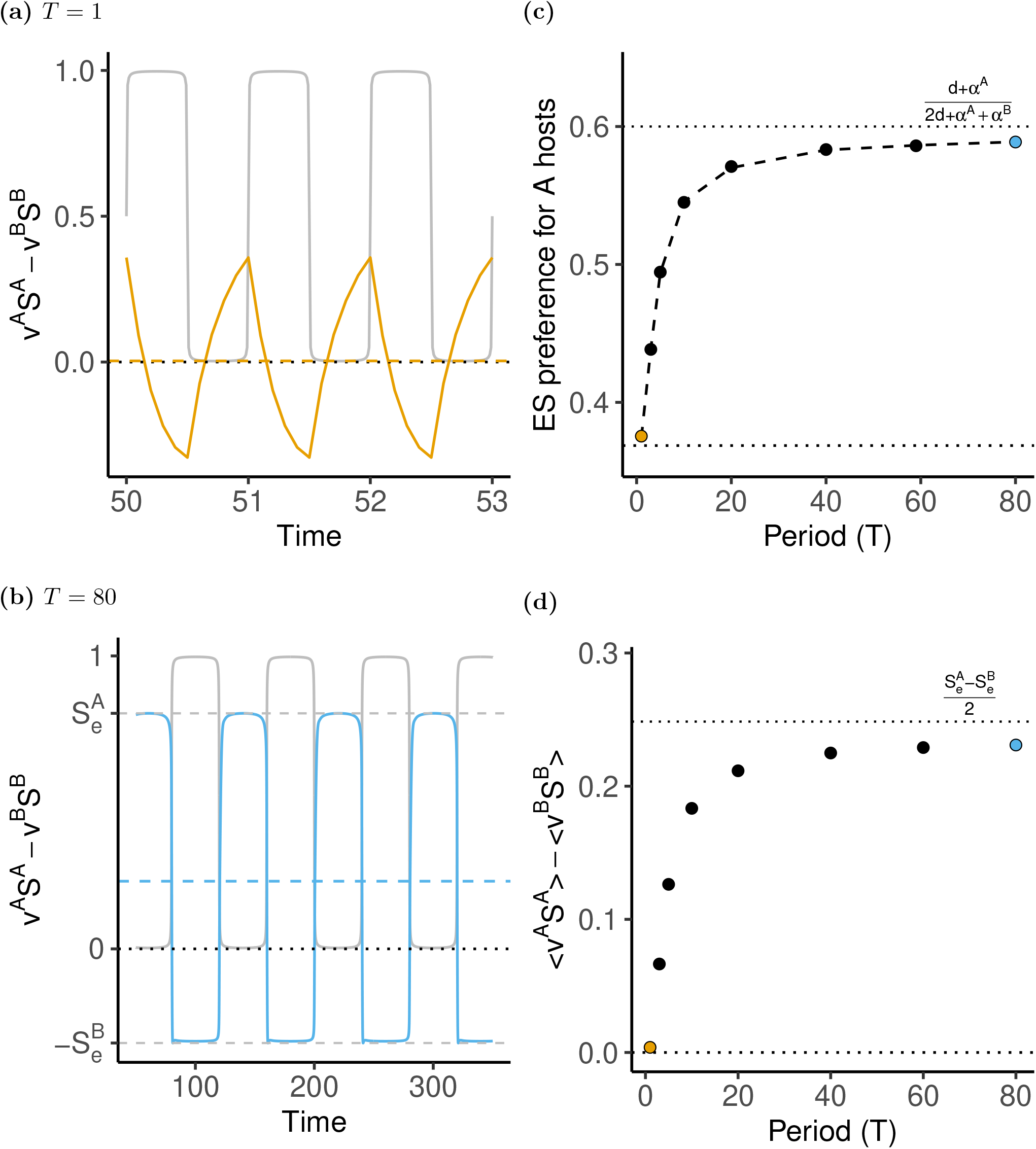
Scenario 2: host preference. (a) Dynamics of *D* = *ν_A_*(*t*)*S_A_*(*t*) – *ν_B_*(*t*)*S_B_*(*t*) for *T* = 1 and *z_w_* = *z_eq_*. The dashed line indicates the mean value of *D*, the dotted line corresponds to the equilibrium value of *D* in the constant environment, which is zero, and the grey line gives the dynamics of *ν*(*t*). (b) Same as (a) but with *T* = 80. (c) Predicted ESS as a function of the period of *ν*(*t*) using the reproductive-value-based approach (dots) and the Floquet exponent (dashed line). The dotted line gives the prediction of the corresponding equilibrium model, *z_eq_* = 0.368652. (d) Mean value of the difference in reproductive output *ν_a_*(*t*)*S_A_*(*t*) – *ν_B_*(*t*)*S_B_*(*t*) as a function of the period for *z_w_* = *z_eq_*. Parameters: *ν*(*t*) = 0.5(1 + (2/*π*) arctan (sin (2*πt*/*T*)/0.01)), *b* = 2, *d^A^* = *d^B^* = 1, *β^A^* = *β^B^* = *β* = 10, *α^A^* = 2, *α^B^* = 1.

**Step 6:** Using the dynamics of the ecological variables, it is possible to calculate the ES strategy for different values of *T* (Appendix S.2) and Figure 5c shows that the ESS increases with the period of fluctuations. For small periods, the ecological variables fluctuate rapidly and the ESS is close to the prediction of the constant model: host preference is biased towards the B hosts because pathogens infecting these hosts have a higher reproductive value. In contrast, when the period of fluctuations is large, host preference is biased towards the A hosts even if their reproductive value is always lower (Figure S2). To understand this counter intuitive result it is important to note that, with the periodic fluctuation we consider, the mean gradient of selection is simply the average over a half-period dominated by host A and a half-period dominated by host B. When the period is large, the epidemiological dynamics reach an endemic equilibrium within each half-period, and this yields:

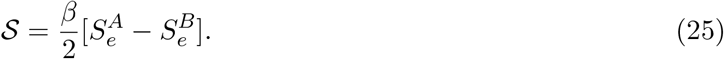

where 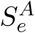 and 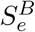 are the densities of the susceptible hosts at the endemic equilibrium of the corresponding single-class models. It is possible to show that 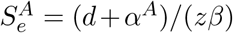 and 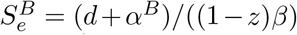. Solving 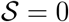 then yields the evolutionary stable strategy *z*\* = (*d* + *α^A^*)/(2*d* + *α^A^* + *α^B^*) which predicts that, indeed, this preference strategy is biased towards host A which suffers more from the infection (i.e., *α^A^* > *α^B^*).

#### Conclusion of Scenario 2

In a constant environment the pathogen evolves a preference for the host which suffers less from the infection because it prolongs the duration of infection. In contrast, we show that slow fluctuations in the abundance of the two hosts can select for the opposite strategy where pathogens evolve a preference for the host that suffers more from the infection. The selection gradient given in equation (23) allows us to shed some light on the effect of temporal fluctuations on pathogen evolution.

### 4.3 Scenario 3: imperfect vaccines

The use of imperfect vaccines may affect the evolution of pathogen virulence and transmission. These evolutionary consequences have been studied by Gandon et al. (2001, 2003) when the coverage of vaccination does not fluctuate in time. In our third epidemiological scenario, we ask how periodic fluctuations in vaccination coverage may affect the evolution of virulence, building on a recent study by Walter & Lion (2021).

We consider the same epidemiological dynamics as in Scenario 2 (equation (21)), but assume that hosts are inoculated at birth with an imperfect vaccine at a rate *ν*(*t*) that fluctuates periodically (figure 3). This yields a fluctuating influx of *A* hosts that are unvaccinated (*θA*(*t*) = *b*(1 – *ν*(*t*))) and *B* hosts that are vaccinated (*θ^B^*(*t*) = *bν*(*t*)).

**Steps 1 and 2:** As in scenario 2, the transition matrix **R***w* is given by equation (22). We now assume that *p_A_* = *p_B_* = 1, and consider that the vaccine can either act by reducing the transmissibility of hosts *B* (*β^B^* = (1 – *r_b_*)*β^A^*) or by decreasing virulence (*α^B^* = (1 – *r_a_*)*α^A^*). This corresponds to the anti-transmission and anti-virulence vaccines introduced by Gandon et al. (2001) (noted *r*_3_ and *r*_4_ respectively in that paper). Finally, we assume that, as in Scenario 1, the trait under selection is the pathogen strategy of host exploitation, *z*, and that transmission and virulence both depend on *z*.

**Step 3:** With these assumptions, the selection gradient at time *t* takes the form

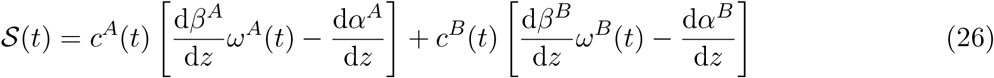

where *c^A^*(*t*) and *c^B^*(*t*) = 1 – *c_A_*(*t*) are the class reproductive values in class *A* and *B* respectively, and

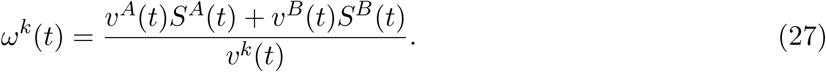

Note that *ω^k^*(*t*) has a useful intepretation: the denominator gives the quality of an “adult” pathogen in class *k*, while the numerator is the expected quality of a pathogen propagule and therefore quantifies the reproductive value of an “offspring” pathogen. So *ω^k^*(*t*) gives a measure of how valuable reproduction is compared to survival in class *k* at any given time.

**Steps 4, 5 and 6:** In general, the densities and reproductive values have complex periodic dynamics (figures S.3 and S.4). However, using the dynamics of reproductive values, it is possible to analytically show (Appendix S.3) that, in the resident population on its periodic attractor:

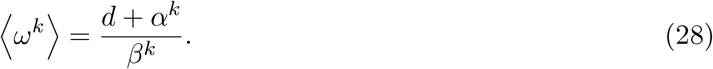

The term on the right-hand-side is one over the basic reproduction ratio *R^k^* of a pathogen when only class *k* is present, and corresponds to the equilibrium value in a model with constant vaccination coverage.

This useful result allows us to rewrite the average selection gradient as

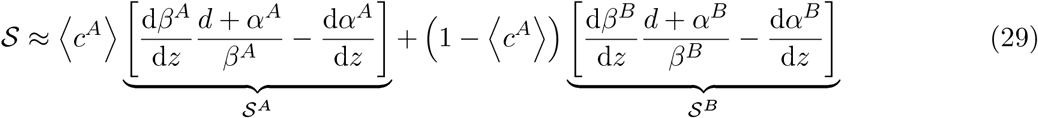

where we have neglected the covariances Cov (*c^k^*,*ω^k^* that arise when taking the mean. Extensive numerical simulations show that this approximation fits very well the prediction of a Floquet analysis (Walter & Lion, 2021).

A full analysis is beyond the scope of this paper, and we refer the reader to the more complete study by Walter & Lion (2021). Nonetheless, is is interesting to see that the selection gradient takes the form of a weighted sum of the selection gradients in class *A* and *B* respectively, exactly as in constant environments. For a constant vaccination coverage, the ES strategy of host exploitation is a weighted mean of the optima in class *A* and in class *B*, respectively given by the zeros of 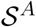 and 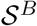. For our anti-transmission vaccine (*r_b_* > 0) 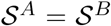, and therefore vaccination has no effect on the optimum. Equation (29) shows that the same holds true for periodic environment, and this is confirmed by numerical calculations of the Floquet exponents (Walter & Lion, 2021). For a vaccine that reduces virulence (*r_a_* > 0), however, 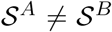, and the position of the ESS is determined by a single variable, which is the mean value, over one period, of the class reproductive value *c^A^*(*t*) in the resident population.

Figure 6b shows that the class reproductive value *c^A^*(*t*) closely tracks the environmental fluctuation *ν*(*t*) when *T* is large (it is close to 1 when *ν*(*t*) = 0, so that only A hosts are produced, and close to zero otherwise), whereas for short periods it quickly fluctuates around its value in a constant environment (figure 6a). An interesting consequence is that, when the period of fluctuations increases, 〈*c^A^*〉 increases (figure 6d), which selects for lower virulence compared to a scenario with constant vaccination coverage (figure 6c). For large periods, 〈*c^A^*〉 converges towards 1/2 (the mean of *ν*(*t*)), which allows the ES virulence to be analytically calculated (Walter & Lion (2021), appendix S.3). Note that, in the latter figure, the slight quantitative discrepancy between the prediction of equation (29) and the Floquet analysis is due to the fact that we have neglected the covariances Cov (*c^k^*,*ω^k^*).

**Figure 6:**
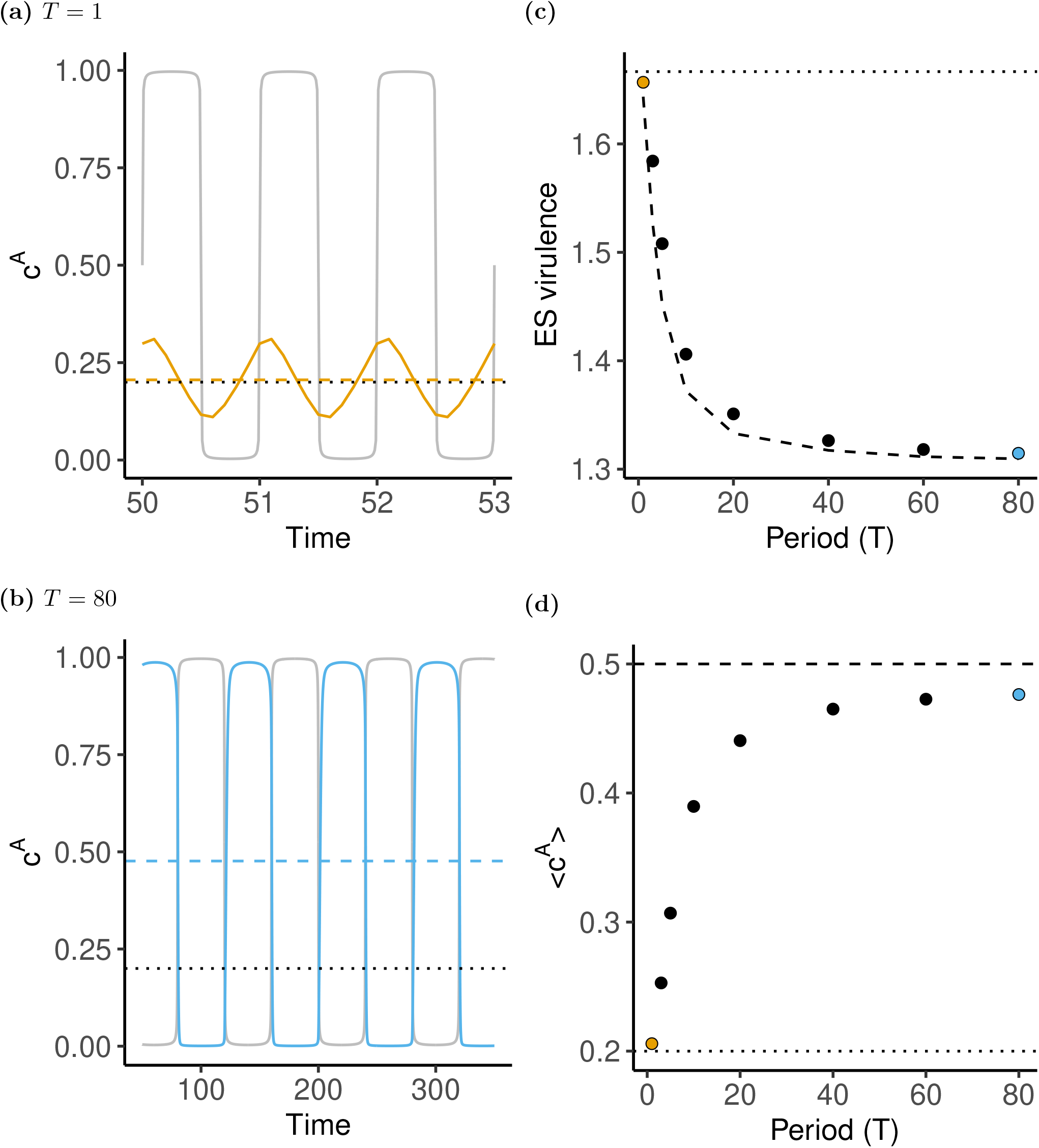
Scenario 3: imperfect vaccines and virulence. (a) Dynamics of the class reproductive value *c^A^*(*t*) for *T* = 1 and *z_w_* = *z_eq_*. The dashed line indicates the mean value, the dotted line the value in a constant environment, and the grey line gives the dynamics of *ν*(*t*). (b) Same as (a) but with *T* = 80. (c) Predicted ESS as a function of the period of *ν*(*t*) using the reproductive-value-based approach (dots) and the Floquet exponent (dashed line). The dotted line gives the prediction of the corresponding equilibrium model, *z_eq_* ≈ 1.667. (d) Mean value of the class reproductive value *c^A^*(*t*) as a function of the period for *z_w_* = *z_eq_*. Parameters: *ν*(*t*) = 0.5(1 + (2/*π*) arctan (sin (2*πt*/*T*)/0.01)), *b* = 2, *d^A^* = *d^B^* = 1, *β^A^*(*z*) = *β^B^*(*z*) = *β* = 10*z*/(1 + *z*), *α^A^* = *z*, *α^B^* = (1 – *r_a_*)*z*, *r_a_* = 0.8.

#### Conclusion of Scenario 3

In a constant environment, vaccination with an imperfect vaccine that affects the within-host growth of the pathogen can select for higher virulence. Here we show that temporal fluctuations in the proportion of vaccinated hosts can mitigate this effect. Importantly, the deviation from the prediction in the absence of fluctuations can be captured by a single quantity, which is the average, over one period, of the class reproductive value *c^A^*(*t*) (see the equation of the selection gradient (29)).

### 5 Discussion

We present a theoretical framework to analyse evolution in periodically fluctuating environments in class-structured populations, and use it to study the evolution of pathogen traits in three epidemiological scenarios. In Scenario 1, we revisit the “Curse of Pharaoh” hypothesis and show that, while propagule longevity is not predicted to affect the evolution of virulence when the environment is constant (Bonhoeffer et al., 1996; Day & Gandon, 2006), fluctuations in the density of susceptible hosts can strongly alter the predictions on the effect of propagule longevity on the evolution of virulence. In Scenario 2, we show that periodic fluctuations in the availability of two types of hosts can bias pathogen preference towards the host where the pathogen has higher virulence, in contrast to the prediction in a temporally constant environment. Finally, in Scenario 3, we show that the evolution of virulence in response to the use of imperfect vaccines is affected by periodic fluctuations in vaccination coverage, which can select for lower virulence compared to constant environments. These three scenarios illustrate how the derivation of the selection gradient (using the general recipe detailed in **Box 1**) can yield important insights on the influence of temporal fluctuations on life-history evolution.

From a methodological standpoint, our analysis extends previous adaptive dynamics studies, which use the Lyapunov (or Floquet, for periodic environments) exponent of a rare mutant as a measure of invasion fitness (Metz et al., 1992; Geritz et al., 1998; Klausmeier, 2008; Metz, 2008; Donnelly et al., 2013; Cornet et al., 2014; Gandon, 2016; Ferris & Best, 2018; Pigeault et al., 2018). Here, we circumvent this problem and derive an expression of the selection gradient on life-history traits, which represents the first-order approximation of invasion fitness for weak selection. We show how it can be used to better understand the impact of environmental fluctuations on the direction of selection and the potential evolutionary endpoints of life-history evolution. In contrast with previous optimisation approaches (e.g. McNamara (1997)), our method does not rely on the assumption that there is no frequency- or density-dependence in the population, and allows us to take into account a variety of ecological feedbacks.

Our approach is based on the analysis of the dynamics of a reproductive-value-weighted frequency of a mutant. This is a recurring idea in evolutionary biology (Fisher, 1930; Taylor & Frank, 1996; Lehmann & Rousset, 2014; Gardner, 2015; Grafen, 2015; Lion, 2018a), but, in contrast to previous approaches (see e.g. Gardner (2014)), the novelty here is that we use a dynamical definition of reproductive value to quantify the fluctuating quality of a class in a periodic environment (see Lion (2018a) for a general discussion on this topic, and Brommer et al. (2000), Caswell (2001), and Bacaër & Abdurahman (2008) for other approaches). The resulting expression of the selection gradient can then be obtained by weighting the effect, at time *t*, of a mutation on the transition rates from class *j* to *k* by the frequency of class *j* at time *t* and by the individual reproductive value of class *k* at time *t*. This has two main implications. First, this allows us to quantify selection at time *t* in terms of the quantity and quality of each class. As in constant environments, the quantity of a class is given by the frequency of individuals in that class, and the quality of a class is measured by their reproductive value, which gives the relative share of future descendants left by individuals in that class. The only difference is that the quantity and quality of each class are allowed to fluctuate periodically over time. Second, the direction of selection can be obtained by computing the average, over one period of the resident attractor, of the instantaneous selection gradient, and we recover a periodic extension of the “invasion implies fixation” principle (Geritz, 2005; Cai & Geritz, 2020; Priklopil & Lehmann, 2020).

It could be argued that, with our approach, the problem of numerically computing a Floquet exponent is replaced by the problem of numerically computing the time-dependent reproductive values and class frequencies. This is of course true if we are simply interested in the quantitative result, but the expression of the selection gradient allows for a qualitative discussion of the selective pressures. The different examples we examine above illustrate that this approach allows us to better understand the often counter-intuitive effects of the periodic fluctuations of the environment on life-history evolution. However, as our approach is currently limited to first-order effects (e.g. convergence stability), the numerical calculation of invasion fitness using Floquet exponents is required to evaluate evolutionary stability and determine whether the predicted evolutionary singularities are ESSs or branching points (Appendix B).

Although periodic environmental fluctuations are important in nature, many organisms also experience stochastic environmental fluctuations. An interesting avenue for future work would be to extend our approach to stochastic stationary ecological dynamics. Since the “invasion implies fixation” principle has also been proven for stochastic fluctuations in the environment (Cai & Geritz, 2020), we think this extension is feasible and would help link our method with classical theory on the influence of environmental stochasticity on life-history evolution (Frank & Slatkin, 1990; Sasaki & Ellner, 1995; Lande et al., 2017). However, this would require a careful definition of the concept of time-dependent reproductive value in stochastic environments.

Our general recipe to study adaptation in periodically fluctuating environments relies on the classical Adaptive Dynamics assumption that the mutation rate is small. This analysis may predict the evolution of generalist strategies that balance the exploitation of the habitats that fluctuate periodically. Yet, as the period of the fluctuation between different environments increases, one may also expect that new mutations will introduce enough genetic variation to allow the population to adapt to this time-varying environment through the recurrent selection of genotypes specialised to each habitat. This would be an interesting avenue for future work.

Although we assumed in our scenarios that all life-history traits are constant, so that temporal fluctuations only come from periodic variations in birth rates which cause fluctuations in densities, it is straightforward to consider time-dependent traits (such as a seasonal transmissibility *β*(*t*)). This would give rise to additional terms capturing the temporal covariance between the trait and some life-cycle-specific ecological variable (Kamo & Sasaki, 2005; Koelle et al., 2005; Cornet et al., 2014; Pigeault et al., 2018).

More generally, fluctuating environments often select for plasticity in life-history traits, which allows an organism to switch between phenotypes specialised to each habitat. It would be interesting to take this into account in our scenarios, especially since there are numerous examples of plastic life-history strategies in pathogens. For instance, malaria parasites have evolved plastic transmission strategies to cope with the fluctuations of their within-host environment as well as the fluctuations of the availability of their mosquito vectors (Mideo & Reece, 2012; Cornet et al., 2014; Birget et al., 2017). In addition, many viruses have the ability to change their host exploitation strategy when they perceive a change in the within-cell environment or other cues from the environment (Gandon, 2016). Our approach can be a valuable tool to understand the evolution of fascinating strategies that enable viruses to coordinate the exploitation of their host population in fluctuating environments (see Bruce et al. (2021) for a recent application of our method).

Our approach provides a general theoretical framework to extend the study of the evolution of life-history plasticity to class-structured life cycles.

## Acknowledgements

We thank P. Carmona, C. Klausmeier and A. Walter for helpful discussions. L. Lehmann, A. Gardner and three anonymous reviewers provided useful comments on previous versions of this manuscript.

## Author contribution

Both authors contributed equally to this work.

## Funding

This work was funded by grants ANR-16-CE35-0012 “STEEP” to SL and ANR-17-CE35-0012 “EVO- MALWILD” to SG from the Agence Nationale de la Recherche.

## Appendix A: Weak-selection approximation of transition rates

To derive the weak selection approximation of the transition rates, we write 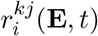 as explicit functions of the phenotypes:

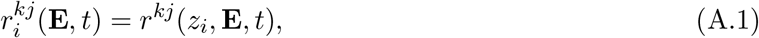

where *z_i_* is the trait of type *i*. Writing *z_m_* = *z_w_* + *ε*, for small *ε*, we use a Taylor expansion to obtain (see e.g. Iwasa et al. (1991), Abrams et al. (1993), Sasaki & Dieckmann (2011), and Lion (2018b))

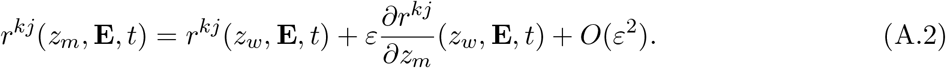

As a result, we have

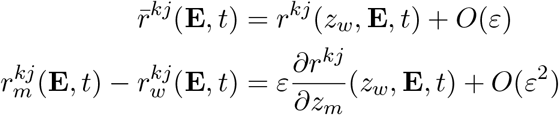

which implies that the dynamics of the class frequencies and individual reproductive values (as well as extrinsic environmental variables) are *O*(1) and functions of 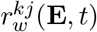 only, while the dynamics of 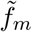 are *O*(*ε*). This leads to a separation of time scales, with the class frequencies and individual reproductive values being fast variables and the weighted mutant frequency 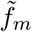 being a slow variable. Importantly, the unweighted average, *f_m_*, is not a slow variable because its dynamics (3) depends on *O*(1) terms (see also Priklopil & Lehmann (2020)).

## Appendix B: The Floquet approach for a rare mutant

The classical Adaptive Dynamics approach is based on the assumption that the mutant is rare and on the derivation of invasion fitness as the Lyapunov exponent of the mutant on the resident attractor (Metz et al., 1992; Geritz et al., 1998; Metz, 2008). For periodic attractors, this amounts to calculating the so-called Floquet exponent of the mutant invasion dynamics (Klausmeier, 2008). Unfortunately, Floquet exponents typically have to be numerically calculated, and only in specific cases can an analytical expression be derived. For instance, when the population has only one class (*K* = 1), it is generally straightforward to derive an analytical expression for the invasion fitness of the mutant by integrating its per-capita growth rate over one period (see e.g. Donnelly et al. (2013) and Ferris & Best (2018)).

The general procedure to calculate the invasion fitness of a rare mutant is to evaluate the mutant dynamics on the resident attractor. This leads to a matrix **R**_*m*_(**E**_*w*_,*t*), where **E**_*w*_ denotes the environment on the resident attractor. One then numerically integrates the matrix differential equation

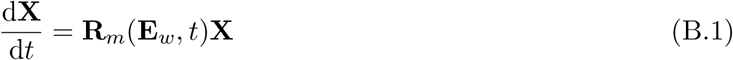

over one period (from *t* = *t*_0_ to *t* = *t*_0_ + *T*) from the initial condition **X**(*t*_0_) = **I**, the identity matrix. The eigenvalues of **X**(*t*_0_ + *T*) are called the Floquet multipliers, and the invasion fitness can then be expressed as the dominant Floquet exponent

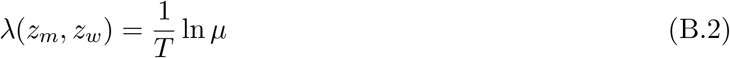

where *μ* is the dominant Floquet multiplier. This numerical procedure can be repeated for any combination of the mutant and resident traits, which makes it possible to develop the full toolbox of Adaptive Dynamics to investigate the convergence and evolutionarily stability of the singularities. Although well established in theory, this method is very rarely encountered in practice in the literature, because very few studies have actually analysed long-term life-history evolution in periodic environments for class-structured populations (but see e.g. Ferris & Best (2018)).

## Appendix S: Supplementary Online Material

### S.1 Scenario 1: Curse of the Pharaoh

Figure S.1a shows that fluctuations in the production of susceptible hosts (top panel) cause fluctuations in the quantity and quality of classes *A* and *B* (middle and bottom panels). By averaging over these fast fluctuations, it is possible to track the frequency *f_m_*(*t*) of the mutant. Figure S.1b shows that our reproductive-value-weighted selection gradient (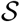, equation (9); dashed line) gives a very accurate prediction for the dynamics of the mutant frequency *f_m_*(*t*) (grey) on the slow time scale (top panel). When one zooms in, we recover the fast fluctuations of the instantaneous selection gradient 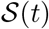 (bottom panel). Note that the direction of selection is well predicted by the sign of 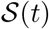.

For the life cycle of Scenario 1, we can use equation (10) to derive the following dynamics of class frequencies:

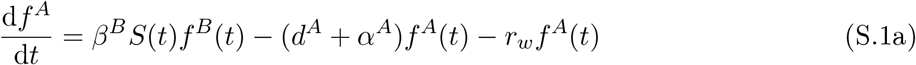

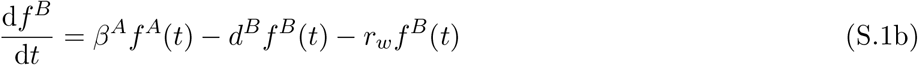

together with the dynamical equation for *S*(*t*). There is no analytical solution in the general case, but numerical integration allows us to investigate the dynamics of the class frequencies. Note that the frequency of free-living pathogens (class *B*) will always tend to lag behind the class *A* of infected hosts because these free-living pathogens are produced from class *A*.

In the absence of fluctuations (*ζ* = 0), the system reaches an equilibrium, which can be calculated by setting d*f^k^*/d*t* and *r_w_* to zero. We thus obtain

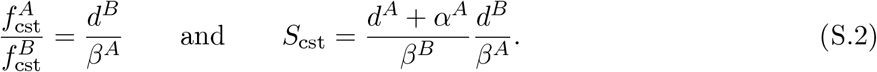

In the presence of fluctuations (*ζ* > 0), an explicit expression of *f^A^*(*t*) on the periodic attractor is beyond our reach, but, because

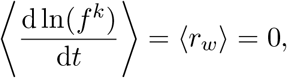

we can show that

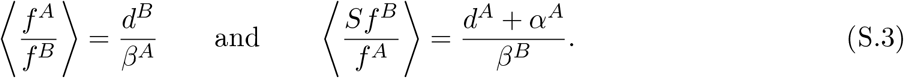

Hence, the ratio of class frequencies, *f^A^*(*t*)/*f^B^*(*t*), fluctuates around an average value *d^B^*/*β^A^*, which corresponds to the ratio of class frequencies in the absence of fluctuations. In particular, this implies that, when *d^B^* → 0 (i.e. when propagules live very long), *f^A^*(*t*) → 0. However, this information cannot be used to further simplify the selection gradient (17).

In contrast, the dynamics of reproductive values yield a very useful expression for the average of the ratio of individual reproductive values, *ν^B^*(*t*)/*ν^A^*(*t*), on the periodic attractor. From equation (12) we obtain

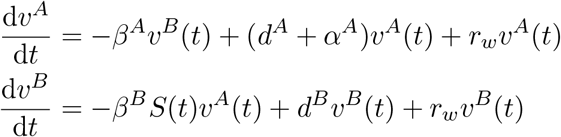

and, using the fact that 〈d ln(*ν^k^*)/*dtE*〉 = 0 on the periodic attractor, we obtain

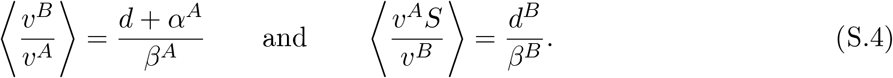

It is straightforward to check that the right-hand sides correspond to the equilibrium values in the absence of fluctuations.

This result suggests that it may be useful to rewrite equation (17) to reveal the selection gradient in a constant environment. We thus write:

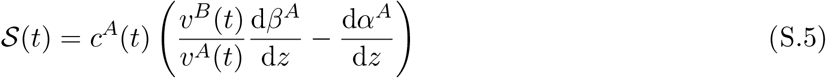

where *c^A^*(*t*) = *ν^A^*(*t*)*f^A^*(*t*) is the class reproductive value at time *t* (Taylor, 1990; Rousset, 2004; Lion, 2018a). Averaging over the period then gives equation (20) in the main text:

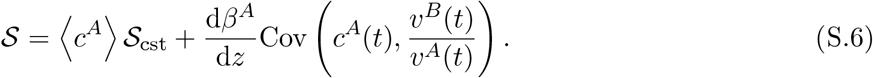

In the absence of fluctuations, or when the covariance is zero, the ESS is predicted from 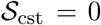, which takes the form of a simple marginal value theorem:

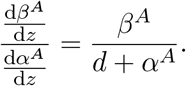

In other words, the ESS maximises the ratio *β^A^*/(*d* + *α^A^*). Any departure from this prediction is caused by a non-zero covariance between *c^A^*(*t*) and 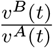. We can develop this covariance and use (S.4) to express this covariance as a function of the reproductive outputs through class *A* (*ν^A^*(*t*)*f^A^*(*t*)) and through class *B* (*ν^B^f^A^*(*t*)). We obtain:

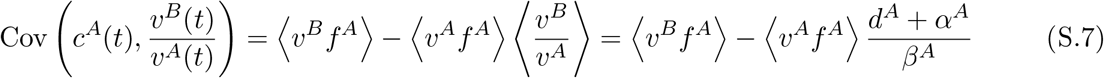

so that the sign of this covariance tells us whether the ratio of the average reproductive outputs is greater or smaller than the average of the ratio of reproductive outputs. In other words, the covariance is positive if

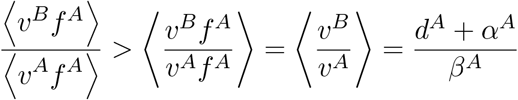

and negative otherwise.

As a final remark on this model, we note that, as in earlier studies (Bonhoeffer et al., 1996; Day & Gandon, 2006), we have neglected the decrease in propagule density due to the infection of new hosts. This effect can be captured by adding a – *β^B^S*(*t*)*I^B^*(*t*) term to the dynamics of *I^B^*, but this has no qualitative impact on our results.

### S.2 Scenario 2: host preference

Figure S.2a shows that, with our choice of periodic fluctuations in *ν*(*t*) (top panel), the system alternates between periods of high densities of susceptible *A* hosts and intermediate densities of susceptible *B* hosts (middle panel). When one host is present, the other is either absent or present at low densities. The resulting dynamics of reproductive values (bottom panel) show that the qualities of *A* and *B* hosts fluctuate over time, but that *B* hosts are always more valuable than *A* hosts. However, some degree of preference for *A* hosts (*z* > 0) may evolve because the densities of susceptible hosts are not equal. This can be shown both for constant and periodic environments.

In the constant case, it is easy to show that, in the resident population at equilibrium

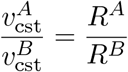

where *R^k^* = *β^k^*/(*d* + *α^k^*) is the basic reproductive ratio in a fully suceptible populations with only *k* hosts present. Thus at the ESS the following condition must be satisfied:

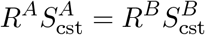

**Figure S.1:**
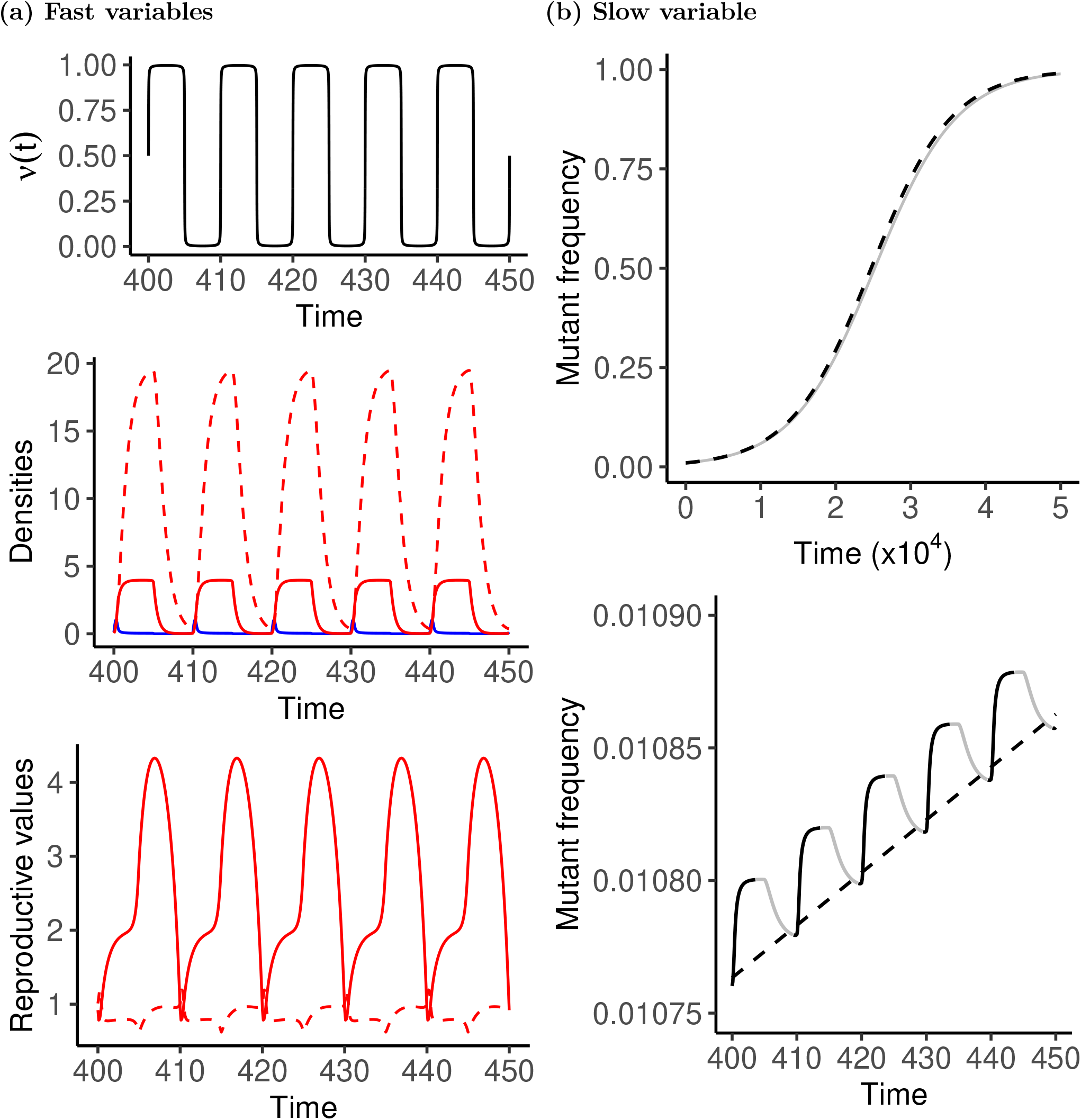
Scenario 1: The Curse of Pharaoh. (a) Ecological dynamics on the fast time scale. Top: periodic probability of production of susceptible hosts, *ν*(*t*). Middle: dynamics of host densities (blue: susceptible hosts, red: infected hosts (solid line: *A* class, dashed line: *B* class (propagules)). Bottom: dynamics of individual reproductive values for pathogens in *A* (solid) and *B* (dashed) hosts. (b) Slow-time dynamics of the frequency of a mutant *f_m_*(*t*) (grey) compared to the prediction using the average selection gradient (dashed line). The lower panel is a zoom showing the oscillations of the mutant frequency on the fast time scale. The direction of selection is well predicted by the sign of the instantaneous selection gradient 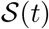 (black: positive, grey: negative). Parameters: *ν*(*t*) = 0.5(1 + (2/*π*) arctan (sin (2*πt*/*T*)/0.01)), *b* = 8, d = *d^A^* = *d^B^* = 1, *β^A^*(*z*) = *β*_0_*z*/(1 + *z*), *β^B^* = *β*_0_ = 10, *α^A^*(*z*) = *z*, *z_w_* = 1, *z_m_* = *z_w_* + 0.005, *T* = 10. The value of *z_w_* is the ESS value in a constant environment with *ν* = 0.5.

which can be numerically solved to yield an intermediate ES value 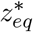.

In the periodic case, the selection gradient vanishes when

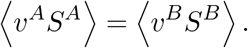

To see how periodic fluctuations can affect the ES preference strategy, we now fix the resident trait at 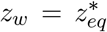, and track the frequency of a mutant with a slightly increased preference for *A* hosts (*z_m_* = *z_w_* + 0.001). In the absence of fluctuations, this mutant should be counter-selected. In figure S.2b, we show that the dynamics of the mutant frequency in the full eco-evolutionary model (solid grey line) is very well predicted, on the slow time scale, by equation (8) with 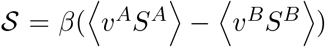 (dashed line), as expected from our general mathematical analysis. Biologically, this means that, although at all times pathogens in *B* hosts have a higher individual reproductive value than pathogens in *A* hosts (figure S.2a, bottom panel), a mutant with increased preference for *A* hosts can still be favoured if the fluctuations in the densities of susceptible hosts tilt the balance in the right direction. In the main text, we show that the deviation from the ESS in the constant environment, 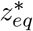, becomes larger as the period, *T*, increases.

In the limit of large periods, for a function *ν* that approaches a step function with minimum 0, maximum 1 and mean 1/2, it is possible to obtain an analytical expression of the ESS. We note that, for large periods, the system approximately behaves as an alternance of single-class equilibria. When only A hosts are present, we have *S^B^* = 0, *ν^A^* = 1 and 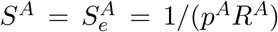, where 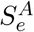 is the equilibrium solution for a population with only A hosts. When only B hosts are present, we have *S^A^* = 0, *ν^B^* = 1 and 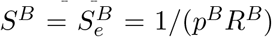, where 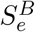 is the equilibrium solution for a population with only B hosts. Thus, we have 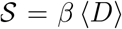 where 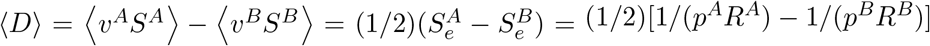, so that, for *p^A^* = *z* = 1 – *p^B^*, the solution of 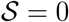 is given by

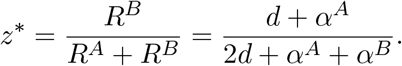

which is the upper limit in figure 5c.

Note that the simulations are performed without any cost of preference, so that the singularity is actually degenerate (i.e. the second isocline of the PIP is vertical near the ESS). However, adding a small constant cost *c* = 1.5(*z* – 1/2)^2^ to the death rates of infected hosts in both classes is enough to make the singularity evolutionarily stable. This only adds a small negative term to the selection gradient that is independent of fluctuations, and has no qualitative impact on our results.

### S.3 Scenario 3: imperfect vaccines

As for scenario 2, figure S.3 shows the dynamics on both the fast, ecological time scale and the slow, evolutionary time scale for a given set of parameter values (chosen such that the trait is at the ESS value in the absence of fluctuations). Again, figure S.3b shows that the averaged selection gradient using reproductive values accurately predicts the dynamics of the mutant frequency *f_m_*(*t*).

The dynamics of the individual reproductive values are given by the system:

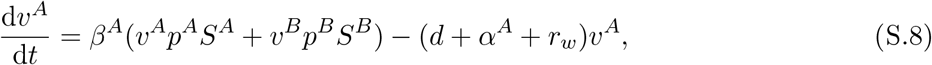

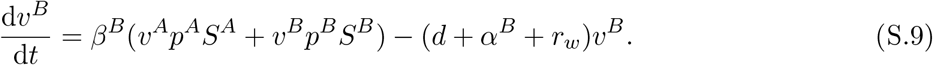

Because on the periodic attractor, we have

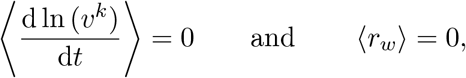

it follows that

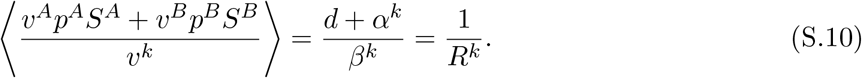

**Figure S.2:**
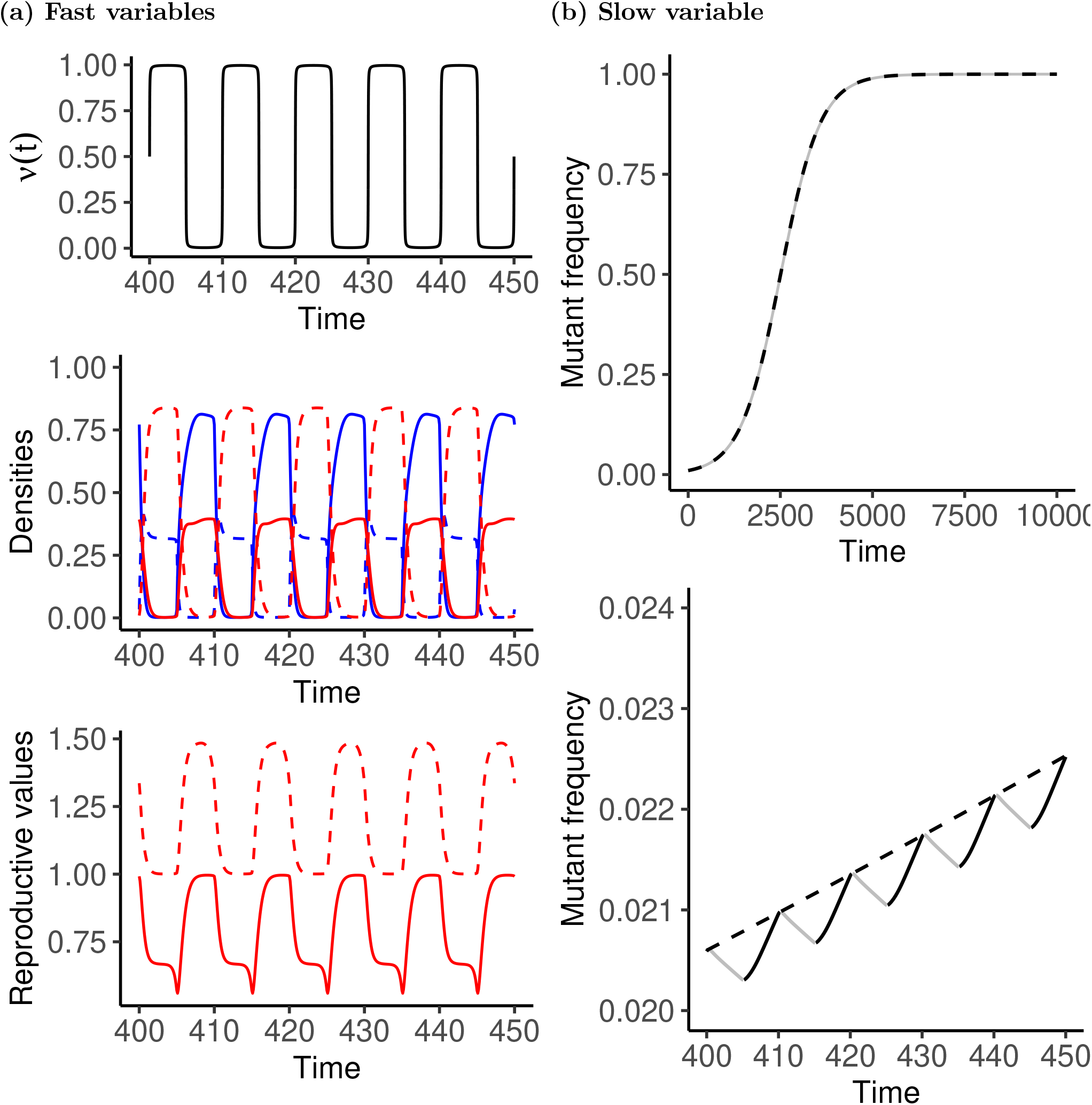
Scenario 2: host preference. (a) Ecological dynamics on the fast time scale. Top: periodic probability of production of *A* susceptible hosts, *ν*(*t*). Middle: dynamics of host densities (blue: susceptible hosts, red: infected hosts, solid lines: *A* hosts, dashed lines: *B* hosts). Bottom: dynamics of individual reproductive values for pathogens in *A* (solid) and *B* (dashed) hosts. (b) Slow-time dynamics of the frequency of a mutant *f_m_*(*t*) (grey) compared to the prediction using the average selection gradient (dashed line). The lower panel is a zoom showing the oscillations of the mutant frequency on the fast time scale. The direction of selection is well predicted by the sign of *ν^A^*(*t*)*S^A^*(*t*) – *ν^B^*(*t*)*S^B^*(*t*) (black: positive, grey: negative). Parameters: *ν*(*t*) = 0.5(1 + (2/*π*) arctan (sin (2*πt*/*T*)/0.01)), *b* = 2, *d^A^* = *d^B^* = 1, *β^A^* = *β^B^* = *β* = 10, *α^A^* = 2, *α^B^* = 1, *z_w_* = 0.368652, *z_m_* = *z_w_* + 0.001, *T* = 10. The value of *z_w_* is the ESS value in a constant environment *ν* = 0.5.

In figure S.4 (bottom panel), we show that the mean of the ratio *ω^k^* = (*ν^A^p^A^S^A^* + *ν^B^p^B^S^B^)/*ν^k^* is indeed equal to 1/*R^k^*. Furthermore, as the period becomes large, *ω^k^* is nearly always equal to 1/*R^k^* except for brief deviations when the environment changes. On the other hand, the middle panel shows that the mean of the reproductive values 〈*ν^k^*〉 are close to their values in a constant environment 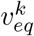 for short periods, but as the period increases, so does the difference between 〈(ν^k^*〉 and 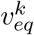.

In the limit of large periods, *c^A^*(*t*) converges towards *ν*(*t*). This can be seen by noting that, for large periods, the model essentially behaves as a succession of single-class equilibria. Half the time, only *A* hosts are present, so that *c^A^* = 1. The rest of the time, there are only *B* hosts and *c^A^* = 0. Hence the mean value of the class reproductive value converges towards 1/2, and the selection gradient simplies to 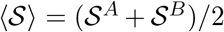. With a trade-off *α^A^* = *z*, *α^B^* = (1 – *r_a_*)*z* and *β^B^* = β^A^ = *β*_0_*z*/(1 + *z*), the ES virulence for large periods has the very simple expression:

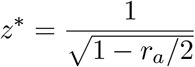

(see also Walter & Lion (2021) for a slightly more general result).

Finally, we note that the dynamics of the individual reproductive values can be used to show that, for a vaccine that linearly reduces transmission (i.e. *r_a_* = 0 and *r_b_* > 0), we have

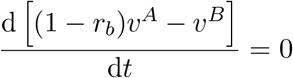

which means that, although the reproductive values fluctuate due to the dynamics of the host densities, their ratio *ν^A^*(*t*)/*ν^B^*(*t*) remains constant and equal to 1/(1 – *r_b_*) at all times. This can be confirmed numerically (results not shown).

**Figure S.3:**
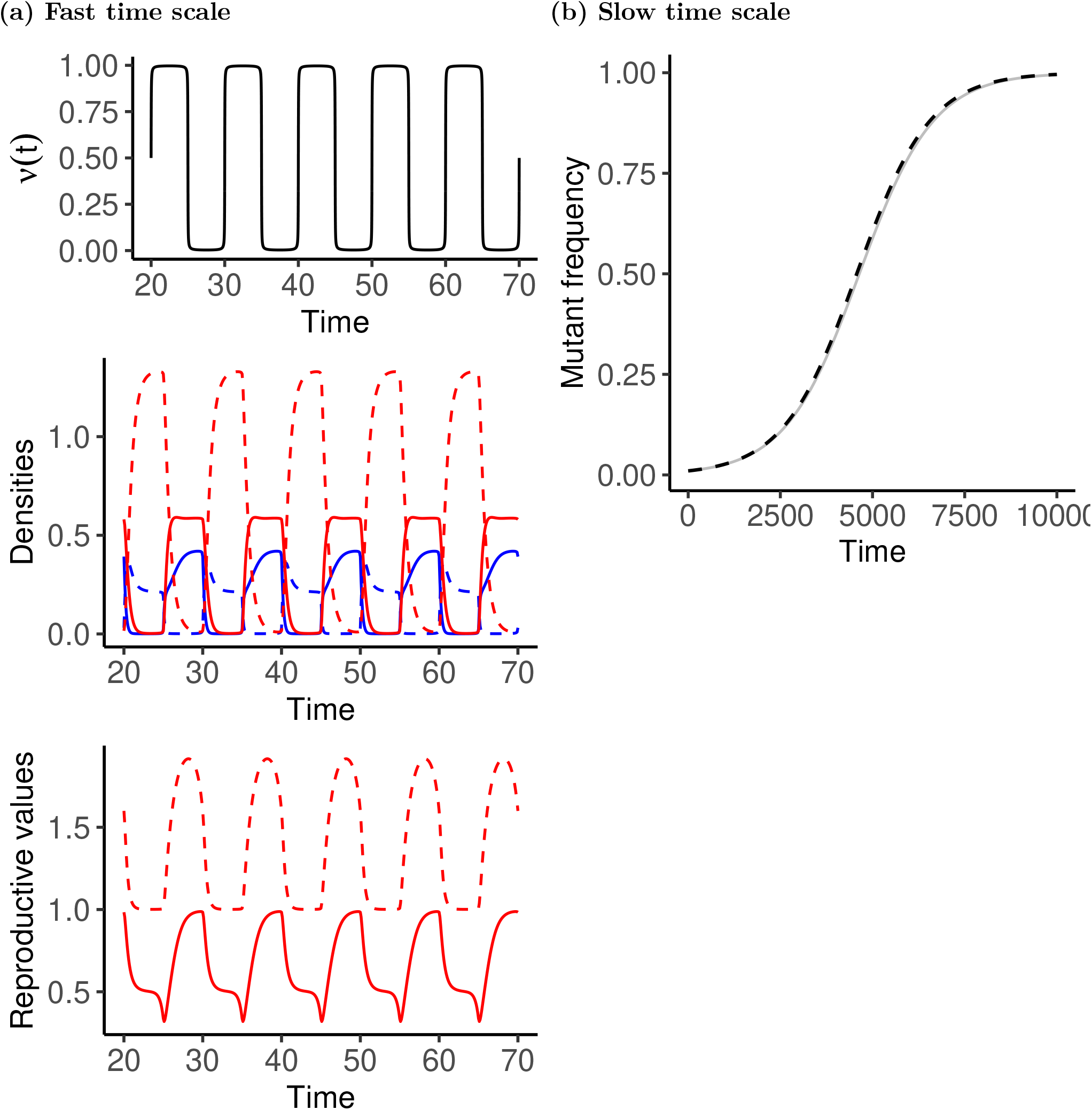
Scenario 3: imperfect vaccine and virulence. (a) Ecological dynamics on the fast time scale. Top: periodic probability of production of *A* susceptible hosts, *ν*(*t*). Middle: dynamics of host densities (blue: susceptible hosts, red: infected hosts, solid lines: *A* hosts, dashed lines: *B* hosts). Bottom: dynamics of individual reproductive values for pathogens in *A* (solid) and *B* (dashed) hosts. (b) Slow-time dynamics of the frequency of a mutant *f_m_*(*t*) (grey) compared to the prediction using the average selection gradient (dashed line). Parameters: *ν*(*t*) = 0.5(1 + (2/*π*) arctan (sin (2*πt*/*T*)/0.01)), *b* = 2, *d^A^* = *d^B^* = 1, *β^A^*(*z*) = *β^B^*(*z*) = *β* = 10*z*/(1 + *z*), *α^A^* = *z*, *α^B^* = (1 – *r_a_*)*z*, *r_a_* = 0.8, *z_w_* = 1.667, *z_m_* = *z_w_* – 0.01, *T* = 10. The value of *z_w_* is the ESS value in a constant environment *ν* = 0.5.

**Figure S.4:**
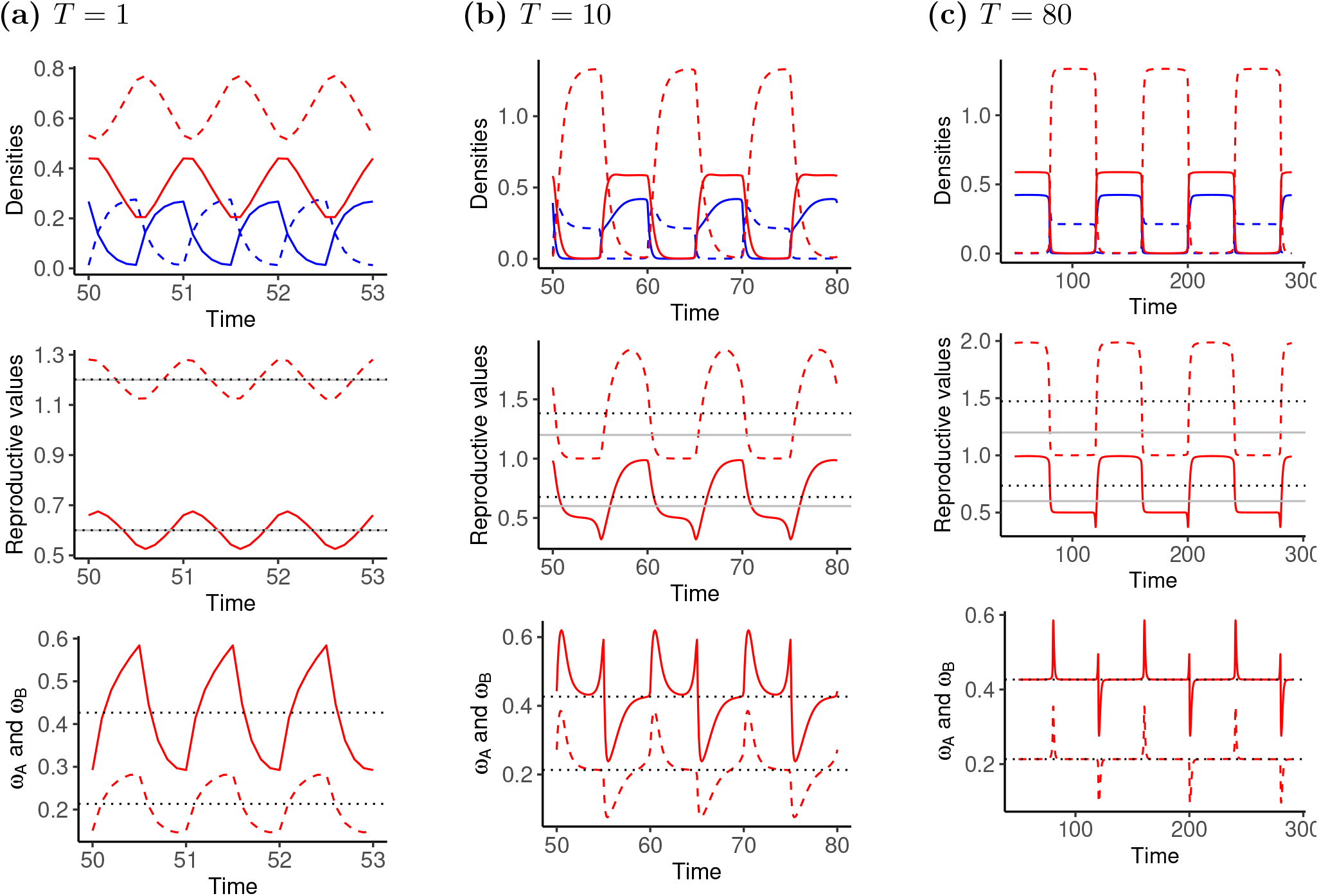
Scenario 3: imperfect vaccine and virulence. (a) Ecological dynamics on the fast time scale for increasing periods. (a) *T* = 1. (b) *T* = 10. (c) *T* = 80. Top: dynamics of host densities (blue: susceptible hosts, red: infected hosts, solid lines: *A* hosts, dashed lines: *B* hosts). Middle: dynamics of individual reproductive values for pathogens in *A* (solid) and *B* (dashed) hosts. The gray and dotted lines indicate the equilibrium and mean values respectively. Bottom: dynamics of *ω^k^* = (*ν^A^S^A^* + *ν^B^S^B^*)/*ν^k^* compared to the mean value (*d* + *α^k^*)/*β^k^*. Parameters: *ν*(*t*) = 0.5(1 + (2/*π*) arctan (sin (2*πt*/*T*)/0.01)), *b* = 2, *d^A^* = *d^B^* = 1, *β^A^*(*z*) = *β^B^*(*z*) = *β* = 10*z/*(1 + *z*), *α^A^* = *z*, *α^B^* = (1 – *r_a_*)*z*, *r_a_* = 0.8, *z_w_* = 1.667. The value of *z_w_* is the ESS value in a constant environment *ν* = 0.5.

## Notes

### Competing Interest Statement

The authors have declared no competing interest.

### Summary of Updates

Minor changes and clarifications.

